# Deep Evolutionary Fitness Inference for Variant Nomination from Directed Evolution

**DOI:** 10.1101/2025.07.22.666175

**Authors:** Max W. Shen, Nathaniel Diamant, Christina Helmling, Raymond Newland, Ziqing Lu, Clara Fannjiang, Simon Kelow, Nathan Frey, Saeed Saremi, Ryan Kelly, Richard Bonneau, Gabriele Scalia, Christian Cunningham, Tommaso Biancalani

**Author notes:** These authors contributed equally to this work (Max W. Shen, Nathaniel Diamant, Christina Helmling, Raymond Newland).

## Abstract

Iterative screening techniques, such as directed evolution, enable high-throughput affinity maturation to optimize binders to molecular interfaces. However, the decision problem of selecting variants from rich, evolved populations to enter low-throughput follow-up methods remains a significant bottleneck. Here, we present evolutionary fitness inference (EVFI) and DeepEVFI, two machine learning methods that model directed evolution from time-series sequencing data, and infer fitness, a variant’s ability to enrich under selection pressure. Our methods flexibly handle mutation mechanisms and starting populations that may be partially unknown – settings relevant to drug discovery – and achieve strong performance on a diverse set of experimental data. We conducted two experimental directed evolution campaigns, using antibodies and macrocyclic peptides libraries to identify and optimize binders to therapeutically relevant targets. EVFI and DeepEVFI identified tighter binders that were missed by human experts using conventional frequency-based approaches, including “rising stars” with low frequency. Beyond initial hit discovery, EVFI and DeepEVFI enables labeling large-scale sequence-fitness datasets and identifying variants of initial binders with diverse properties.

## 1 Introduction

Directed evolution and iterated screening assays such as yeast surface display, phage display, and mRNA display, are powerful methods for protein engineering and drug discovery, in which a diverse population is evolved through multiple rounds of selection, replication, and potentially mutation [1–4]. The ability to link desired biological activity to survival rate enables the engineering of biomolecules for a wide variety of properties [5, 6]. *In vitro* display assays can explore substantial sequence space over time, with library sizes up to 10^9^ in yeast, 10^11^ in phage, and 10^14^ for mRNA display [2–4, 7, 8]. Mutations during directed evolution can arise from many sources, are difficult to completely eliminate, and improve the exploration of fitness landscapes [9, 10]. Timepoint populations from directed evolution can contain vast genotypic diversity with rich dynamics of competing variants emerging, flourishing, and declining over each round of evolution [11].

Computational modeling of time-series sequencing data from directed evolution experiments can serve many purposes, such as illuminating sequence-fitness relationships [11–18], and mining the rich sequence diversity in evolved populations for hit finding [10, 11, 13, 15, 16]. A particular noteworthy purpose, for drug discovery, is addressing the decision problem of variant nomination which entails how a researcher should follow up from a directed evolution campaign to pick a small number of variants, dozens to hundreds, for functional measurement assays which are typically expensive and low-throughput (**Figure 1a**). In drug discovery, evolved populations may contain promising leads with strong binding affinities or beneficial biochemical properties, but suboptimal variant nomination strategies can mean failure to discover these variants. This decision problem of variant nomination thus plays a key role in campaign success. In the yeast, phage, and mRNA display literature, variants are often nominated by sampling the final population and ranking each identified binder by overall frequency. While this method does identify promising candidate binders, it is often suboptimal as it does not truly leverage the large-scale data that directed evolution offers [4–8, 19–22]. Instead, fitness inference – inferring each variant’s fitness, or quantifying its ability to survive and grow under selection pressure and competition – can enable the discovery of more promising variants that are less biased by nuisance factors (**Figure 1b**) [11, 12, 16, 18, 23]. A rich body of work has developed computational modeling methods when exactly two timepoints are available, such as for deep mutational scanning [16–18, 24], comparing positive to negative sorts [23, 25], and methods that use deep learning [21, 22, 26].

**Fig. 1.**
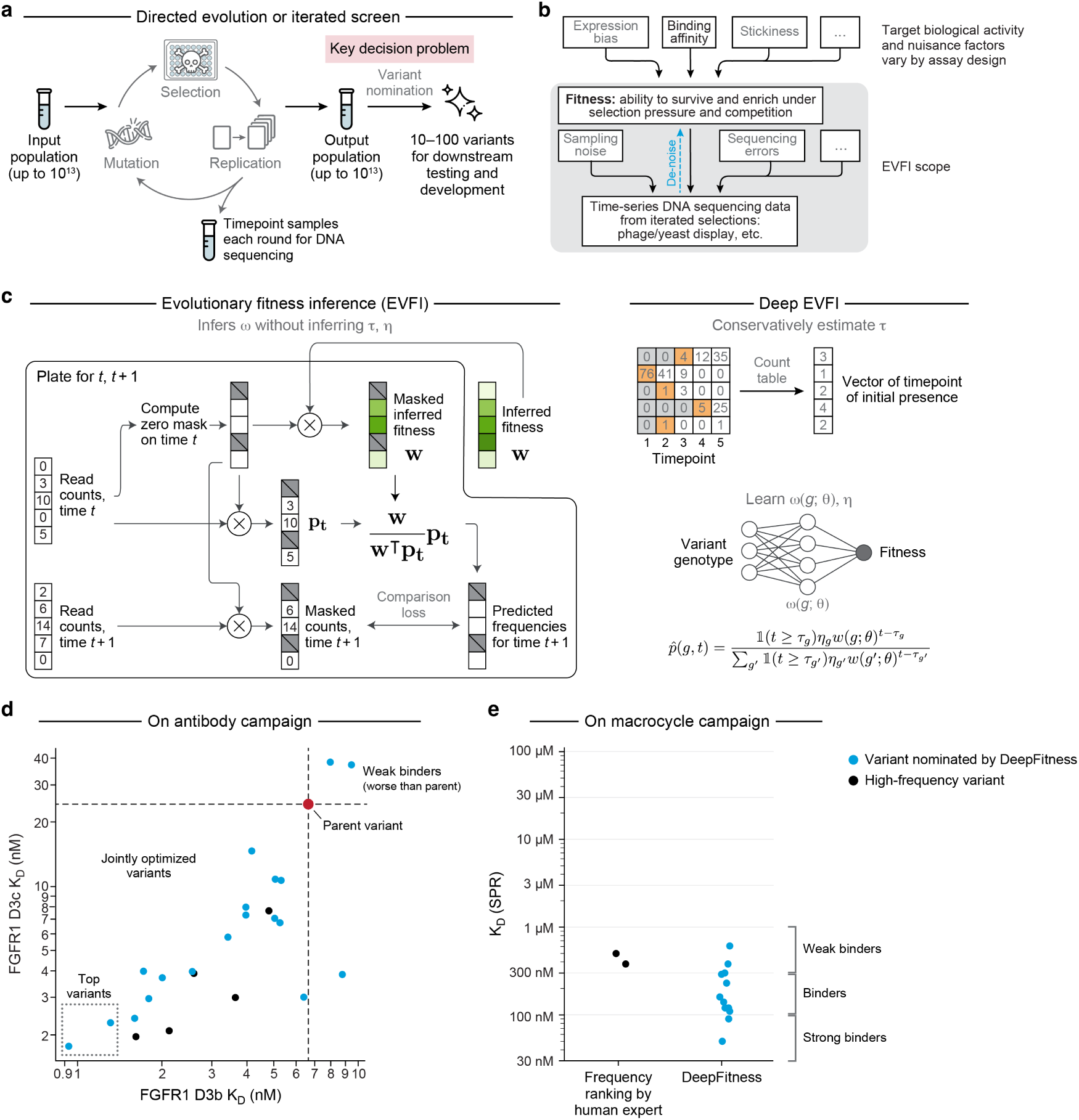
EVFI and DeepEVFI are machine learning methods for fitness inference. **a**. Schematic of the decision problem of variant nomination for directed evolution or iterated screens. **b**. Conceptual schematic of the relationship between binding affinity and fitness. **c**. EVFI (left) and DeepEVFI. Symbols and variables are defined in the results, methods, and supplement. **d**. Experimental results on antibody screen (also reported in Figure 4c). EVFI nominates two top variants which outperform high-frequency variants **e**. Experimental results on macrocycle screen (also reported in Figure 5b). DeepEVFI identifies multiple binders that outperform the two most enriched peptides at the end of the campaign.

In contrast, there is a relative lack of computational methods designed to support conditions of directed evolution relevant to drug development, which often involve three or more timepoints, partially known or unknown mutation mechanisms between selection rounds, and highly diverse, poorly characterized starting populations [2, 3, 5, 6, 11]. Existing methods that support three or more timepoints include ACIDES, Enrich2, Rosace, and AMaLa, but they make restrictive assumptions on mutations between rounds, or starting populations [12, 16, 24, 27]. As higher mutation rates, more selection rounds, and more diverse starting populations can enable broader exploration of fitness landscapes to find better variants, these settings are powerful and relevant for drug discovery applications.

Here, we present EVFI and DeepEVFI, two machine learning methods for fitness inference tailored to realistic iterative drug screening scenarios, including *in vitro* directed evolution experiments like yeast, phage, and mRNA display. EVFI and Deep-EVFI infer variant fitness from time-series DNA sequencing data of variant frequencies using a temporal dynamics model, without relying on low-throughput, expensive functional measurements like binding affinity (**Figure 1c**). In contrast to prior methods, EVFI and DeepEVFI are i) designed for datasets with two or more timepoints, ii) do not require specifying a particular model of mutation, while supporting data with or without mutations, or with unknown mutation mechanisms, and iii) can model directed evolution starting from a single wildtype sequence, or from diverse initial populations that are not fully characterized. By introducing EVFI and DeepEVFI as separate methods, we distinguish the contributions from using deep learning from other methodological contributions.

We study a variety of public datasets and find that EVFI is best-in-class among methods that do not use deep learning, while DeepEVFI uses deep learning to further improve performance. Using fitness inference, we find that “rising star” [11] variants with high inferred fitness but low frequency, are common in publicly available mRNA, phage, and yeast display datasets, which suggests that fitness inference can improve variant nomination outcomes in practice.

To investigate the effectiveness of EVFI and DeepEVFI in directed evolution experiments, we conducted two campaigns on two different therapeutic modalities, antibodies and macrocycles, and compared our inferred fitness methods to the more traditional frequency-counting method to identify and optimize binders against therapeutically relevant targets. In our first campaign, we optimized an antibody to simultaneously bind to two closely related variants of FGFR1, a key oncology target [28–30], called D3b and D3c. EVFI identified the top two variants, with the highest showing 7.1× and 13.8× improved binding to D3b and D3c, respectively (**Figure 1d**), over the parental variant. For the thioether-linked peptide macrocycle campaign targeting the YAP domain of TEAD2 (also a key oncology target [31–34]), we used DeepEVFI to identify potent 8-mer macrocycle binders by selecting sequences with high inferred fitness and low experimental frequency previously not identified with traditional analysis methods (**Figure 1e**). We achieved a 90% hit rate, with 92% of DeepEVFI binders matching or exceeding the affinity of the best human-selected binder. The best DeepEVFI binder had 7.6× improved affinity compared to the best human-selected binder. Overall, EVFI and DeepEVFI successfully nominated the tightest binders in both campaigns as validated by surface plasmon resonance (SPR).

## 2 Results

### 2.1 Evolutionary fitness inference from time-series data

We consider the problem of fitness inference from *T* timepoints of frequency measurements of a population of *G* unique variants undergoing iterated rounds of selection, replication, and possibly mutation. We use a standard population genetics definition for fitness [35]

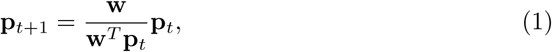

where **p***_t_* is a vector of variant frequencies at round *t*, and **w** is a vector of variant fitness values. By this definition, fitness quantifies a variant’s ability to survive and grow under selection pressure and competition. For hit finding in drug discovery, selections are designed so that improved binding affinity to a target protein is one of the primary drivers of selection survival, but overall fitness may also be influenced by other factors, such as non-productive binding (e.g. hydrophobicity), amplification biases, and expression variability (**Figure 1b**).

We briefly highlight a core methodological contribution here – how we address fitness inference with flexibility to mutation mechanisms – and provide a mathematical foundation of fitness inference and further details in the **Methods** and **Supplementary Information**.

Mutations introduce new variants at low frequencies that may not be present in DNA sequencing data, which introduces optimization challenges when using Eq. (1). To achieve flexibility for different models of mutation, EVFI and DeepEVFI are built on a data generative process that considers the round *τ* when each variant first enters the population, and each variant’s initial abundance *η* (**Supplementary Information**). Inferring fitness then requires jointly inferring ***w***, ***τ***, ***η*** which is challenging, so other methods have introduced restrictive assumptions to make the problem more tractable. Here, instead of introducing restrictive assumptions, we instead use estimate certain latent variables in a data-driven manner. EVFI infers fitness using a masked optimization approach based on the presence of zero counts in consecutive timepoint pairs (**Figure 1c**, left), which is equivalent to using conservative data-driven estimates for ***τ*** and ***η***. DeepEVFI jointly learns a sequence-to-fitness neural network for fitness inference and learns ***η***, using a conservative data-driven estimate of ***τ*** (**Figure 1c**, right; **Methods**), which we show improves inference for variants in the training set, evaluated on held-out selection rounds. Both methods model temporal dynamics with Eq. (1) by using it to simulate populations forward in time based on inferred fitness values. Simulated populations are compared to observed populations and a loss is calculated based on the data likelihood under a noise model accounting for genetic drift and sampling noise. The loss is backpropagated to update inferred fitness values or the sequence-to-fitness neural net.

Flexibly handling mutations is important as mutations play a key role in powerful directed evolution platforms like yeast, phage, and mRNA display. Mutations can occur at roughly 10^−3^ mutations per construct (m/c) per timepoint in phage and mRNA display, and 10^−6^ m/c per timepoint in yeast display (**Supplementary Information**). Furthermore, mutations arise from diverse sources, can be difficult to fully anticipate or eliminate, and are often desirable for finding better optima in the sequence-fitness landscape. With its increased flexibility, our methods are applicable to a wide variety of data, including data with mutation mechanisms that are difficult to precisely characterize, partially or completely unknown, and data where the existence or absence of mutations is unclear.

### 2.2 EVFI and DeepEVFI improve predictions on future selections

We studied seven public datasets of time-series DNA sequencing data of populations undergoing multiple selection rounds, including optimization of human protein domains, antibodies, adeno-associated virus (AAV), snoRNAs, and peptides, using methods including phage display, yeast two-hybrid, yeast growth, and *in vivo* directed evolution for AAV transduction [36–41] (**Figure 2a**; **Extended Data Table 1**). These datasets were previously studied using the ACIDES method [12] and are denoted by the letters A-G. We also studied two new datasets: a yeast surface display campaign identifying antibodies for FGFR1 binding with six timepoints (denoted Y-Ab), and an mRNA display campaign identifying macrocyclic peptides for binding to TEAD with seven timepoints (M-MP). All datasets and their results are used in this section to compare the predictive power of EVFI and DeepEVFI against existing methods but are discussed in more detail in subsequent sections.

**Fig. 2.**
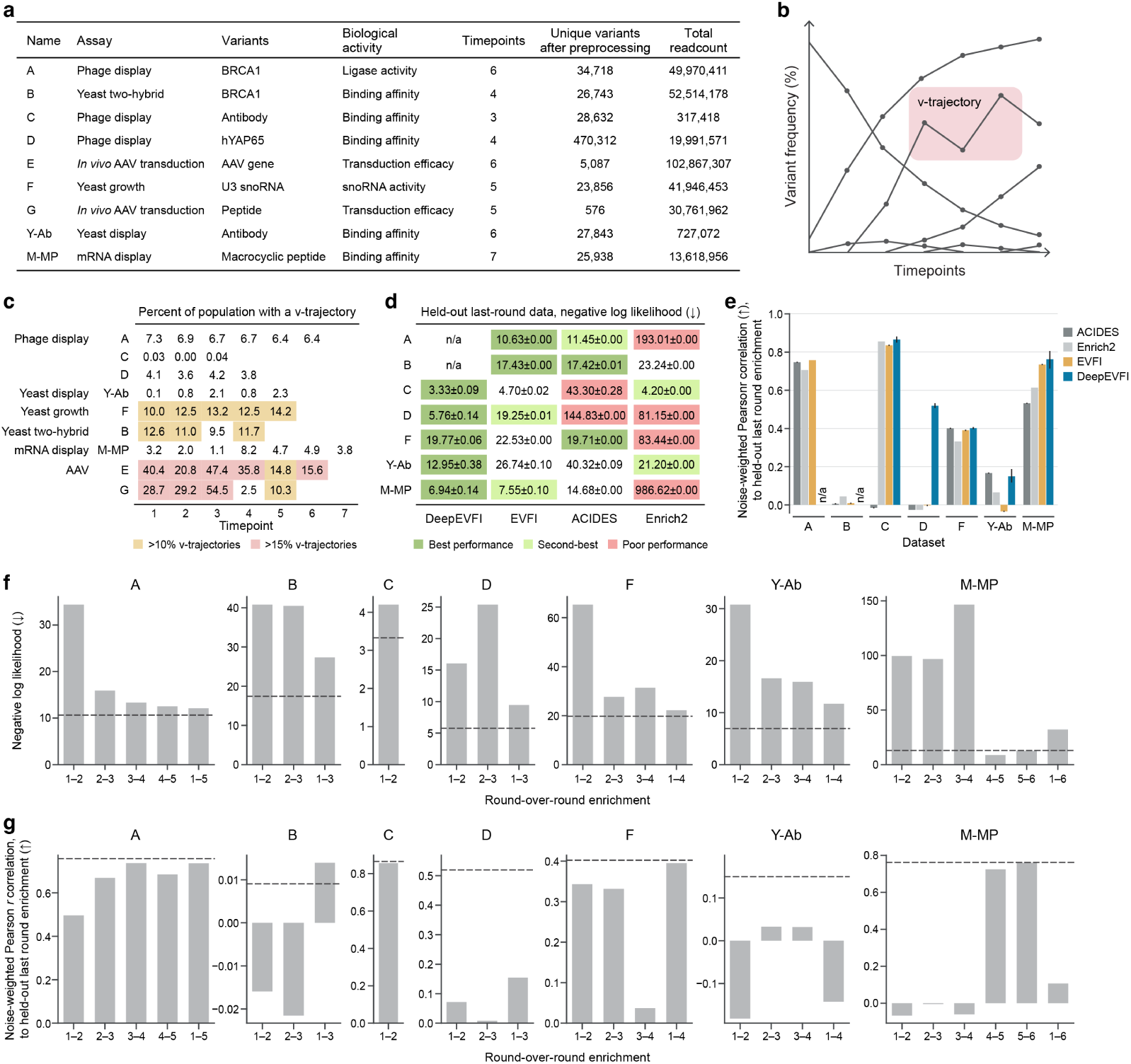
EVFI and DeepEVFI exhibit state-of-the-art performance on prediction of future selections. **a**. Description of nine datasets studied in this work. **b**. Example of v-trajectory. **c**. Percent of timepoint populations comprised of variants with a v-trajectory at any timepoint trio, with high confidence (>95% probability). Yellow shows *>* 10%, red shows *>* 15%, indicating stronger violations of model assumptions. **d**. Multinomial negative log likelihood (NLL; bits per variant; lower value is better) comparing predicted final timepoint variant frequencies to held-out data, reported as mean plus-or-minus standard deviation over five training replicates. Dark green shows best performing method(s) within one standard deviation. Light green shows second-best, and red shows significantly poor performance. **e**. Noise-weighted Pearson correlation between predicted final timepoint variant enrichment to held-out data. Bars and error bars plot mean and standard deviation over five training replicates. **f** -g. NLL and noise-weighted Pearson correlation of round-over-round enrichment compared to final round enrichment, as in d-e. Blue line indicates EVFI or DeepEVFI.

We sought to compare our method to ACIDES [12] and Enrich2 [16], which are representative existing methods for modeling enrichment over three or more timepoints [24]. Both of these methods, as well as ours, assume that variants have a time-invariant growth rate. However, prior work has not determined whether real-world datasets conform to these model assumptions. To investigate this, we developed a statistical measure to assess whether, and to what extent, datasets violate these assumptions (**Methods**). We mathematically prove that, under these assumptions, a “v-trajectory” — a timepoint triplet where a variant decreases and then increases in frequency, forming a “V” shape (**Figure 2b**) — cannot occur (Supplementary Information). To detect the presence of v-trajectories in real datasets, we computed the probability that triplets of observed counts indicate a v-trajectory under binomial sampling noise model, and scored the datasets using the total frequency of variants with a v-trajectory with high confidence (*>*95% probability; **Methods**). This analysis provided a principled framework for deciding whether fitness inference methods can be meaningfully applied to a real-world dataset.

In the nine datasets that we studied, we found that the two *in vivo* AAV datasets [39, 41] exhibit significant violations with up to 47.4% and 54.5% of the population containing v-trajectories (**Figure 2c**). In contrast, yeast growth and yeast two-hybrid campaigns exhibit mild v-trajectory violations (ranging 10% — 15%), while yeast, phage, and mRNA display campaigns showed *<* 10%. Therefore, we excluded *in vivo* AAV selection datasets E and G, leaving seven datasets for further analysis.

For each remaining dataset, we trained each method using data up to the second-to-last timepoint and used the final timepoint as a holdout test set. We evaluated the ability to predict variant frequencies at the final timepoint for variants that are also present at the second-to-last timepoint. This approach focuses the evaluation on enrichment modeling instead of predicting which novel variants enter the population. We compared EVFI to ACIDES and Enrich2, as all of these methods do not use deep learning, and do not use genotype information (**Figure 2d**). EVFI performed best, achieving 2.1× improved likelihood than ACIDES (geometric mean across datasets) and 4.5× improved likelihood over Enrich2. Additionally, EVFI achieved the best likelihood on four out of seven datasets and otherwise performed closely to the best method. We observed that ACIDES and Enrich2 performed well on some datasets, but poorly on others, such as on dataset A (10.63 for EVFI, *vs.* 193.01 for Enrich2; lower is better) and on dataset D (19.25 for EVFI, *vs.* 144.83 for ACIDES). In the Supplementary Material, we analyze potential reasons why ACIDES and Enrich2 can perform poorly. Taken together, EVFI performed strongest on these seven datasets, with reliable performance across a variety of iterated selection campaigns spanning phage display, yeast growth, yeast display, and mRNA display.

Next, we evaluated DeepEVFI which uses genotype information as an input for deep learning, unlike ACIDES, Enrich2, and EVFI. Our evaluation was restricted to five out of the seven datasets where genotype data was available. DeepEVFI achieved the strongest likelihood performance, improving 1.6× over EVFI, 4.6× over ACIDES, and 7.1× over Enrich2 (geometric mean across datasets). We computed noise-weighted Pearson correlation to compare the predicted and observed final-timepoint enrichment which downweights low-count variants with higher noise (**Figure 2e**). In all five datasets, DeepEVFI achieves the strongest noise-weighted correlation to held-out enrichment or ties with the best performing method within one standard deviation across training replicates.

A key property of our problem setting is the availability of three or more timepoints. However, it is common to perform enrichment analysis using only two timepoints. We evaluated our methods against two-round enrichment methods that compute fitness by dividing variant frequencies in an “after” round by a “before” round. A complication of this approach is how to choose which two timepoints, among *T* (*T* − 1) pairs, to use.

We find that performance varies widely by which timepoints are used to compute the two-round enrichment and observed that no single choice is best for all seven datasets. EVFI and DeepEVFI match or exceed the best performance of this commonly used round-over-round enrichment calculation using either consecutive timepoint pairs or last-over-first (**Figure 2f-g**). By using all timepoint data, fitness inference bypasses the timepoint choice required for two-round enrichment analysis and achieves better performance.

### 2.3 Rising stars are common and overlooked

A common approach for variant nomination is sampling variants from the final timepoint population. For fitness inference to improve on this, there must exist higher fitness variants unlikely to be sampled from the final timepoint due to low frequency, which we term “rising stars“, a concept previously studied in the continuous evolution setting [11]. However, the extent to which rising stars exist in yeast, phage, and mRNA display in the context of drug discovery has received little systematic investigation.

We investigated evidence for rising stars by comparing EVFI inferred fitness to final timepoint frequencies across the seven datasets (**Extended Data Figure 1**). In all seven datasets, we find dozens to thousands of variants with evidence of having high fitness yet lower frequency (**Figure 3**), than *baseline variants* which we take as the highest frequency variant in the final timepoint. Formally, we define rising stars as variants with over 90% probability of having higher fitness than the baseline variant under a Dirichlet-Multinomial model to account for noise and fitness uncertainty from low read counts (**Methods**).

**Fig. 3.**
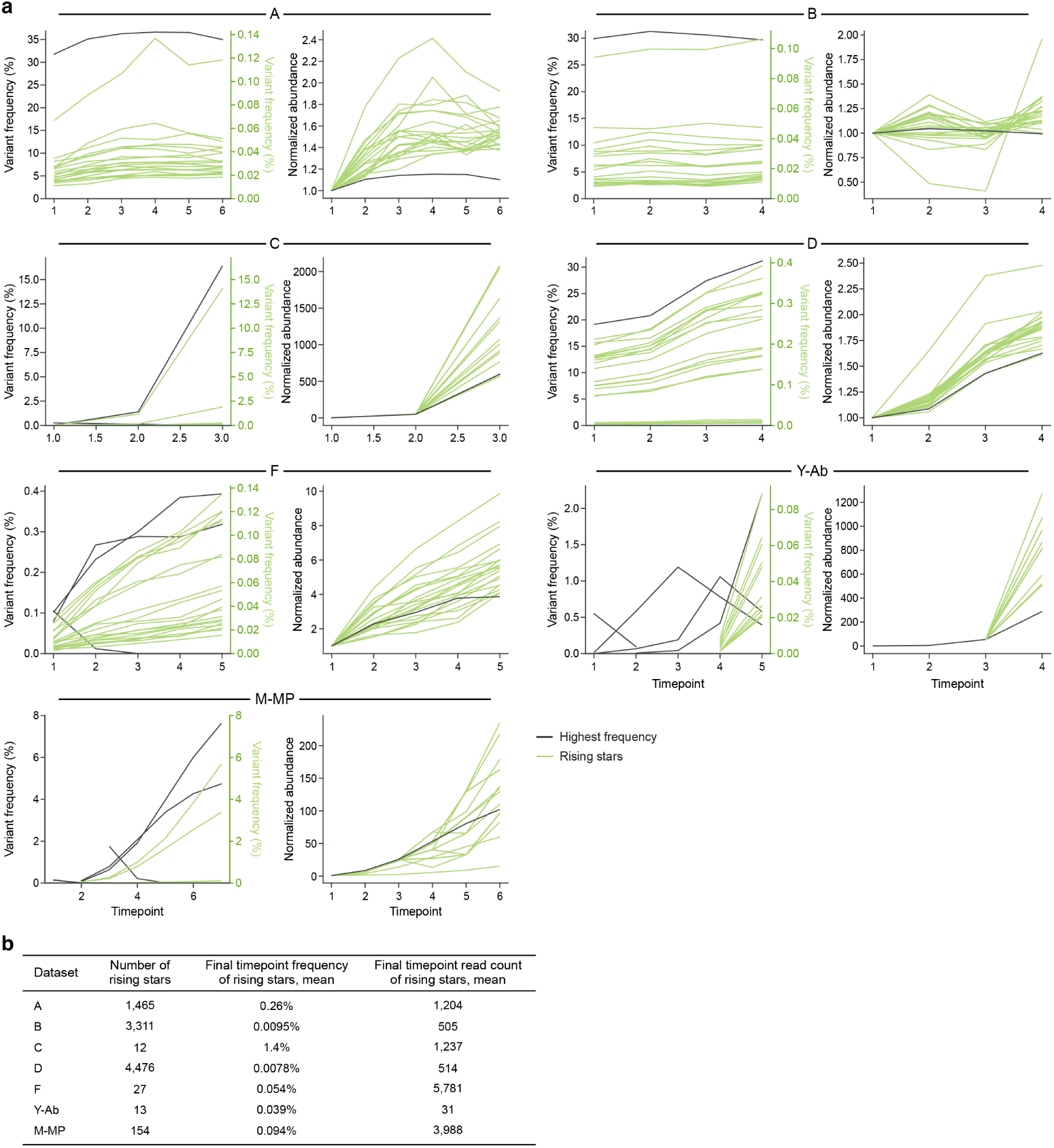
Rising stars are common across real world datasets. **a**. Variant frequency trajectories. Black shows variants with highest frequency in a timepoint. Green shows rising stars. Normalized abundance trajectories are plotted by computationally adjusting all initial variant frequencies to be equal. **b**. Statistics of rising star variants in datasets.

Overall, rising stars had 182× lower final frequency than baseline variants (geometric mean across the seven datasets). In the M-MP dataset, 154 rising stars had a final timepoint frequency averaging 0.094%, approximately 100× lower than the baseline at 7.6%. Importantly, we saw clear read-count and trajectory evidence supporting that rising stars have strong enrichment and cannot be explained solely by measurement noise. Rising stars had an average of 3,988 sequencing reads in the final timepoint in the M-MP dataset, 31 reads in dataset Y-Ab, and hundreds to thousands of reads in datasets A-D and F. Visual inspection of rising star trajectories confirmed observed evidence of consistent enrichment outcompeting the baseline over multiple timepoints (**Figure 3a**).

Empirically in these datasets high final timepoint frequency was largely determined by high initial frequency and less so by fitness, which is problematic when initial frequency is determined by factors unrelated to the biological activity of interest. Rising stars had 10-100× lower initial frequencies than the baseline variant which preventing them from overtaking the baseline within the number of selections performed. To investigate what could have happened if rising stars had the same initial frequency as the baseline variant, we computationally adjusted initial frequencies in the dataset while preserving inferred growth rates, and simulated evolutionary dynamics using equation 1. We found that rising stars would have dominated the baseline in final-timepoint frequency (**Figure 3a**).

Rising stars are common and abundant across seven diverse datasets, including phage display, yeast growth, yeast display, and mRNA display, consistent with similar findings in continuous directed evolution platforms [11]. These results suggest that sampling from the final timepoint population can be suboptimal for identifying the highest fitness variants and demonstrate that fitness inference can identify rising stars with stronger growth under selection pressure and competition.

### 2.4 EVFI enables the optimization of a sub-nanomolar antibody using fewer selection rounds

We next applied EVFI to the affinity optimization of 1C2, a previously isolated singlechain variable fragment (scFv) antibody directed against two closely related variants of fibroblast growth factor receptor 1 (FGFR1) called D3b and D3c. FGFR1 is a receptor tyrosine kinase involved in cell growth, differentiation, angiogenesis, and tissue repair. FGFR1 antagonists, such as monoclonal antibodies and peptide inhibitors, are in clinical trials for their potential to improve cancer treatment outcomes by inhibiting oncogenic signaling, angiogenesis, and overcome resistance to existing therapies [28–30]. The parent antibody has an affinity of 6.6 nM affinity to D3b and 24.2 nM to D3c. We constructed a library diversifying all six complementarity determining regions (CDRs) and subjected the population to five rounds of selection, utilizing section conditions ranging 250 nM target concentration to as low as 100 pM (**Methods**).

A human expert scientist sampled clones from populations after four rounds as well as after five rounds of selection to nominate variants for testing. In total, 71 clones were sampled yielding 25 unique variants. All variants were successfully expressed as full length hIgG1s and SPR was used to measure their binding affinities to D3b (**Figure 4a**, black dots). Additionally, five of the tightest D3b binders were manually chosen to measure their D3c binding affinity (**Figure 4b**, black dots).

**Fig. 4.**
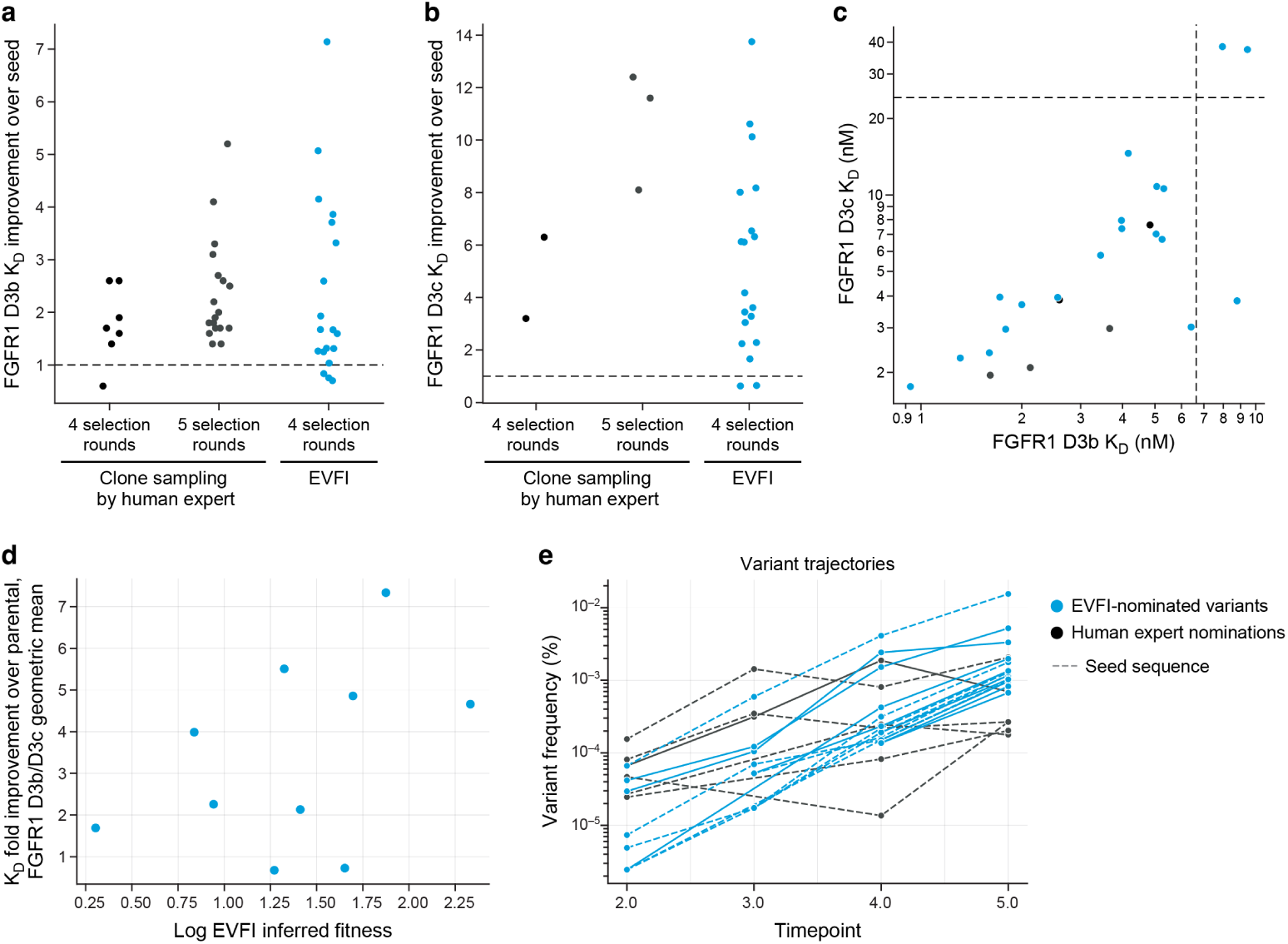
EVFI identifies a sub-nanomolar antibody without using all selections. **a-c**. Binding affinities measured through dissociation constant *KD*, for D3b (**a**), D3c (**b**), and both (**c**). Dashed gray line indicates parent variant. **d**. Comparison of log inferred fitness of nominated truncated variants *vs.* the geometric mean of *KD* fold improvement on binding D3b and D3c over parent variant of full-length nominated variants. Spearman *ρ*=0.51, *p*=0.06, *N* =14. **e**. Frequency trajectories of variants. In all panels, blue depicts EVFI-nominated variants using four selection rounds, gray depicts human expert nominations by sampling clones from the final timepoint after four or five selection rounds.

Subsequently, we tested whether EVFI could nominate tight binders using fewer selection rounds than were used previously. As the complete heavy and light chain sequences were too long for Illumina DNA sequencing read lengths, we only read the full heavy chain and a portion of the light chain sequence with Illumina DNA sequencing to identify what we call “truncated variant” sequences. We then used EVFI to infer fitness from the first four selection rounds and excluded the data from the fifth selection round. From this dataset, we nominated truncated variants and extended them with the parental light chain to obtain full-length antibody variants for testing. Among the 25 variants identified with EVFI, 20 were able to be expressed and 19 showed confirmed binding in SPR as hIgG1 (**Extended Data** Figure 2).

To understand how EVFI compared to traditional analysis techniques, we compared the binding affinities to D3b and D3c in three groups: human-expert-nominated variants after four selection rounds (H-4S); human-expert-nominated variants after five selection rounds (H-5S); and EVFI-nominated variants after four selection rounds (EVFI-4S).

Overall, the EVFI-4S group had tighter binders than H-4S, and found comparable binders to H-5S. For D3b affinity, none (0/7) of the H-4S group improved the parent variant affinity by more than 3× while 32% (6/19) of EVFI-4S achieved affinities did. A similar rate was observed in the H-5S group at 22% (4/18) (**Figure 4a**). For D3c affinity, no H-4S variants improved the parent variant affinity by more than 10×, while two H-5S variants did and three EVFI-4S did (**Figure 4b**). Many EVFI-4S variants (37%; 7/19) had equal or better affinity to D3b and D3c than the best H-4S binder.

The best overall dual-binding variant was nominated in EVFI-4S which showed an improved affinity against D3b at 7.1× (930 pM) and D3c at 13.8× (1.76 nM) (Figure 4c). Notably, EVFI-4S variants experienced weaker selection pressure (target concentration ≥ 1 nM) than H-5S variants (≥ 100 pM). Collectively, these results demonstrate that EVFI identified higher affinity binders than clone sampling given the same rounds of selection. Additionally EVFI was able to discover comparable binders earlier than the clone sampling reducing resources and discovery timelines for similar results.

Among EVFI-nominated variants, inferred fitness scores positively correlated with combined D3b and D3c affinity improvement over parent variant (*ρ*=0.51, *p*=0.06, *N* =14; Spearman correlation) (**Figure 4c**). EVFI-4S variants demonstrate strong and consistent enrichment over multiple timepoints compared to H-4S variants (**Figure 4e**), which have inconsistent and weaker enrichment trajectories. EVFI-4S variants had an average frequency of 0.25% after four selection rounds. Overall, these results demonstrate that EVFI variant nomination can perform similarly, and in some cases exceed, sampling from the population after performing an additional selection round, even at weaker selection stringency. EVFI may thus enable identifying strong variants earlier with reduced experimental effort.

### 2.5 DeepEVFI identifies macrocyclic peptides with high binding affinity

We performed two mRNA display campaigns with the goal of identifying thioether macrocyclic peptides binders against the YAP binding domain of TEAD2. TEAD proteins are transcription factors that regulate the downstream effects of the Hippo signaling pathway that controls organ size and development [31–34]. TEAD serves as a receptor for its co-activators YAP/TAZ to upregulate cell proliferation and survival and the enhanced activity of YAP/TAZ and TEAD has been observed in various human cancers [42–45]. Macrocyclic peptides have proven highly effective for engaging previously-deemed “undruggable” shallow grooves and clefts observed in many proteinprotein interactions such as TEAD-YAP.

Given TEAD transcription factors are intracellularly localized we designed small cyclic lariat libraries with 8-9 amino acids as lower molecular weight scaffolds may have a better potential for passive membrane permeability. However, lower molecular weight peptide libraries have traditionally been difficult to discover in directed evolution experiments due their loss in the wash steps prior to elution as a result of their general lower affinity and fast binding kinetics.

After 7 rounds of selection, the final timepoint of campaign 1 was sequenced and 21 hit peptides were nominated by a human expert using frequency ranking for affinity determination. Twenty of the hit peptides were also synthesized with GKK tags at the C-terminus in order to improve their solubility [46], resulting in 41 total hit peptides. Twelve of the synthesized 41 variants showed binding in SPR (**Figure 5b**), with affinities ranging from 1-100 *µ*M.

**Fig. 5.**
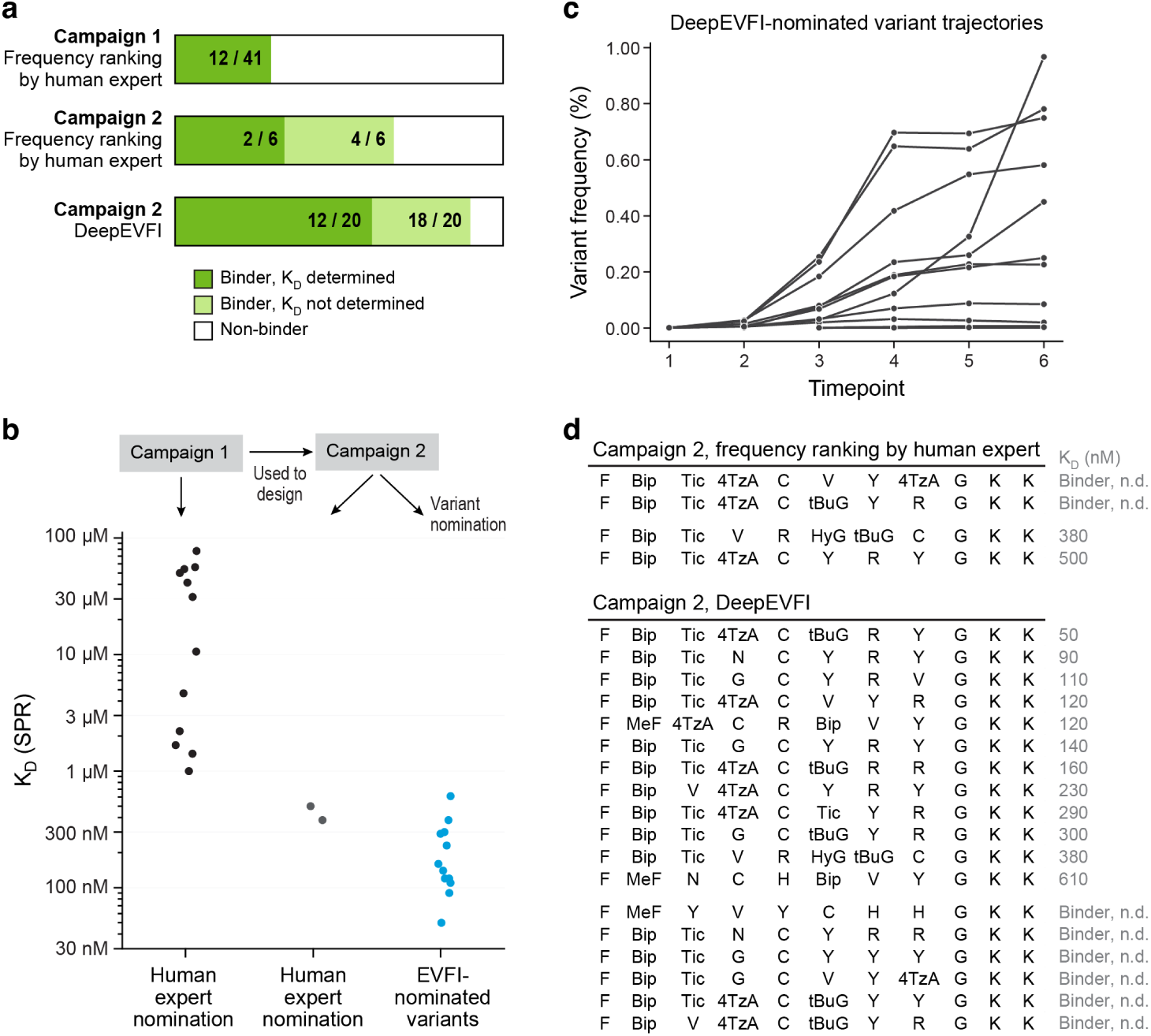
DeepEVFI identifies tighter binders than frequency-based nomination in a short macrocycle directed evolution campaign. **a**. Diagram depicting SPR outcomes for nominated variant sets on macrocyclic peptides binding to TEAD. b. TEAD *KD* for variants with determined SPR values. c. Frequency trajectories of DeepEVFI-nominated variants in M-MP dataset. d. Genotypes and SPR binding affinity.

From these results, a second selection campaign was designed based on the first campaign using only 8mer lariats and incorporating only one wash step instead of three in each round of selection to help prevent their loss before the elution step. After 7 rounds of selection, a human expert manually nominated six hit peptides using frequency ranking from the sequencing data of the final timepoint population. As was done in the first selection, each hit peptide was synthesized with a -GKK tag on the C-terminus to help with solubility during affinity measurements. Of the six variants, two did not bind, two had measurable binding affinity but their *K_D_* could not be determined, and two were binders with *K_D_*of 380 nM and 500 nM.

We applied DeepEVFI on the DNA sequencing data of campaign 2 by jointly inferring fitness and learning a sequence-to-fitness deep neural network. We nominated a set of variants using inferred fitness, while also designing for variant diversity (**Methods**), without knowledge of the six human-expert nominated variants, which yielded one duplicate renomination (380 nM binder).

Among DeepEVFI-nominated variants, 90% (18/20) bound, and affinity was determined for 12 variants (**Figure 5a**; **Extended Data** Figure 3). Nearly all DeepEVFI variants (92%; 11/12) matched or improved binding affinity over the best human-expert nominated variant, which DeepEVFI also renominated. The single best discovered binder had 50 nM affinity, which is 7.6× tighter than the best human-nominated variant from campaign 2, and 20× tighter than the best from campaign 1 (**Figure 5b**). Notably, DeepEVFI-nominated variants had an average frequency of 0.2% in the final timepoint (**Figure 5c**), and the tightest binder had a frequency of 0.02% in the final timepoint. Our discovery of an 8-mer macrocycle with 50 nM affinity to TEAD approaches the 15 nM affinity of a previously reported 17-mer macrocycle [47], despite being shorter (**Figure 5d**).

Overall, these results demonstrate that DeepEVFI using deep DNA sequencing of all timepoints from a directed evolution campaign can nominate variants with tighter binding affinity that are overlooked by common variant nomination strategies like frequency ranking.

### 2.6 Variant nomination with diversity

Discovering diverse hits is crucial for lead discovery in drug development. A lead candidate must optimize multiple objectives and property constraints, though multiobjective optimization problems do not inherently care about diversity beyond the variation in drug candidates on the Pareto frontier balancing trade-offs between competing objectives. A more direct motivation for diversity is solving multi-objective optimization problems when not all objectives or property constraints are known ahead of time, which is common in practice.

In the seven datasets we studied, despite undergoing two to seven rounds of selection, none of the populations converged to a small group of dominant variants. Instead, they maintained high diversity, with thousands to tens of thousands of unique variants, and frequencies typically followed an exponential distribution (**Extended Data** Figure 4).

Indeed, visual inspection of UMAP plots from different datasets reveals that variants with high final timepoint frequencies cover limited sequence diversity, while variants with high inferred log fitness span a much broader range of sequence diversity within the population (**Figure 6a-b**). This led to the hypothesis that fitness inference could be used to nominate diverse variant sets with high inferred fitness, reflecting evidence of consistent strong enrichment over timepoints, which may better represent the large sequence diversity present in evolved populations.

**Fig. 6.**
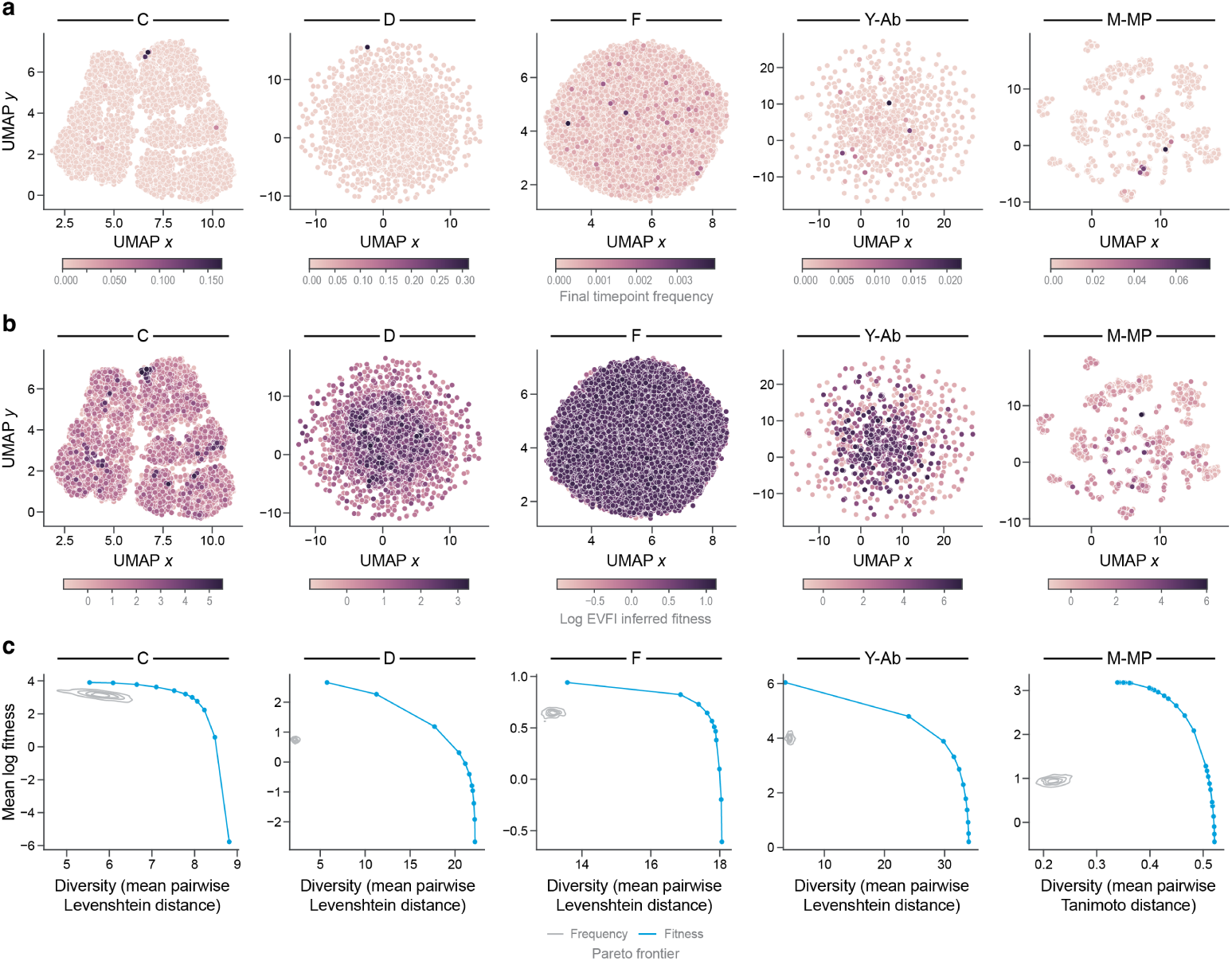
Diversity of variant sets nominated by final timepoint frequency vs. inferred fitness. a-b. UMAP plots of sequences, colored by final timepoint frequency (a) and log inferred fitness (b). c. Pareto frontier of nomination sets with 96 variants optimizing average log fitness or average pairwise distance (blue) compared to nomination sets by sampling 96 variants by final timepoint frequency (gray).

Across the five datasets with genotype information, we find that final timepoint sampling produces variant sets with substantially lower diversity and lower fitness than variant sets we build using full time-series DNA sequencing data (**Figure 6c**). To build these variant sets, we used varying trade-offs between high inferred log fitness and high sequence diversity using a greedy algorithm (**Methods**), which in all cases yields variant sets with higher mean fitness, higher diversity, or both, compared to final timepoint sampling.

We note that in datasets D and Y-Ab, final timepoint sampling nominates variants sets with less than five point mutation difference between variant pairs on average, while we build nomination sets that differ by more than 20 mutations from each other on average in dataset D, and more than 30 mutations in Y-Ab. In datasets Y-Ab and M-MP, our top fitness-optimized variant sets have 7.4× higher fitness than variant sets from final timepoint sampling. Taken together, fitness inference also supports nominating diverse variants that can better capture the sequence diversity in evolved populations.

## 3 Discussion

In this work, we present EVFI and DeepEVFI, which are intentionally designed with a high degree of flexibility, support two or more timepoints, allow mutation mechanisms that are unknown or only partially known, and enable analysis of starting populations that are diverse and may only be partially characterized. This flexibility contrasts with previously developed methods many of which are limited to exactly two timepoints or are limited to specific assumptions on mutation mechanisms or starting populations. This flexibility underpins EVFI and DeepEVFI’s strong performance on a broad range of public datasets, and makes EVFI and DeepEVFI highly relevant methods for drug discovery, in which a variety of assays are commonly used, such as yeast, mRNA, and phage display.

We empirically reported that rising stars are common in many public directed evolution datasets, spanning a variety of assays and optimized biomolecules. This observation suggests that many directed evolution assays are not near convergence and ndicate that sequencing-based fitness inference methods like EVFI and DeepEVFI may be broadly useful and provide value to many directed evolution campaigns in practice.

Using yeast display with antibodies and mRNA display with macrocyclic peptides, we demonstrate that EVFI and DeepEVFI enable the identification of tight binders with low frequency that may easily be missed by conventional nomination approaches ike clonal sampling and final timepoint frequency ranking. Beyond hit discovery, finding diverse variants with comparable or improved affinity compared to human expert picks EVFI and DeepEVFI supports hit expansion which may assist lead discovery. Large sequence-fitness datasets may shed light on sequence-activity relationships, assist researchers in future designs and follow-up studies such as testing mutation reversions, and provide alternative leads for development towards the clinic.

We remark that fitness inference is based on variant frequencies in time-series sequencing data and thus are based primarily on measured changes in frequency. Inferred fitness therefore corresponds to a variant’s ability to survive and grow under selection pressure, but may have an imperfect relationship with a specific desired biological activity such as binding affinity even if in principle the selection is designed primarily on binding affinity. However, if enrichment is caused more by nuisance factors than by the primary activity of interest such as binding affinity EVFI or DeepEVFI may be less useful for nominating improved variants. EVFI thus benefits from careful design of selection conditions. Toward that end, EVFI can also be used as a framework for quantifying selection design quality by considering the correlation between denoised enrichment over multiple timepoints with measurements of the primary activity of interest. It thus may be possible to use EVFI to improve selection quality, which in turn improves the utility of EVFI.

## 4 Methods

### 4.1 Yeast display on FGFR1

The CDR-H3 from clone 1C2 was subsequently incorporated into a new scFv YSD ibrary with a diversity of 2*x*10^8^, introducing non-germline encoded diversity in CDR-L1, CDR-L3, CDR-H1, and CDR-H2. The library was adapted into pCTCON2 and transformed by electroporation. This library formed a starting population we call A. Selection was performed iteratively with splitting some populations and subjecting each split to different selection conditions, to yield a total of seven final populations, with a total of 15 intermediate populations A-O. The seven tracks yielding each final population is: 1) ABCDE, 2) ABCDFJ, 3) ABCDFK, 4) ABCDGL, 5) ABCDHM, 6) ABCDHN, 7) ABCDIO. The selection conditions yielding each population is B: MACS against 125 nM D3b and 125 nM D3b in 500 microliter beads; C: FACS on 10 nM D3b and 10 nM D3c; D, E: 10 nM D3b/D3c; F: 1 nM D3b/D3c; G/L: 100 pM D3c; H: 1 nM D3b/D3c with 1-hr FACS buffer wash; I/O: 1 nM D3c with 1-hr FACS buffer wash with 1 µM untagged competition; J: 1 nM D3b; K: 1 nM D3c; M: 1 nM D3b with 1-hr FACS buffer wash; N: 1 nM D3c with 1-hr FACS buffer wash. Clones were sampled from all final populations, which were E with four prior selection rounds, and J, K, L, M, N, and O with five prior selection rounds. Sampled clones were reformatted into human IgG1 and analyzed by SPR. Timepoint populations after each selection were also subjected to next-generation sequencing (NGS) using the Illumina 2×300 MiSeq platform. Given the 600 base pair limit of this method, each output was amplified from the Light Chain Framework 3 through the entire Heavy Chain. Sequencing data were then analyzed using EVFI and DeepEVFI as described in this paper. Selected designs from this analysis were expressed, and SPR was performed.

### 4.2 FGFR1 Naive Yeast Display Library Construction & Seed Antibody Isolation

A single-chain variable fragment (scFv) library was designed in a VL-linker-VH format and synthesized (Twist Bioscience). The design incorporated diversity predominantly within the CDR-H3 region while the remaining CDR’s (Complementary Determining Regions) were germline encoded. The library utilized germline scaffolds heavily prevalent in the human repertoire and previous clinical antibodies. Sequences coding for the overlap region with the pCTCON2 vector were appended as appropriate to the scFv design allowing for homologous recombination in yeast (See Supplementary Data for overlap sequences). The pCTCON2 vector was prepared for homologous recombination by digestion overnight with SalI-HF followed by digestion overnight with NheI-HF and BamIHF (New England Biolabs). The library was amplified via PCR using Q5 HotStart High-Fidelity Polymerase (New England Biolabs). The amplified scFv library insert and the digested vector were transformed into RJY100 yeast strain[48], using previously described electroporation methods[49, 50]. The final library contained 1.6*x*10^9^ members.

Sorts for FGFR1-D3b & FGFR1-D3c binders were conducted using previously described yeast surface display techniques[49, 50]. Initial rounds of sorting were conducted using Magnetic Activated Cell Sorting (MACS) via AutoMACS (Miltenyi Biotec), ensuring at least ten fold coverage of the library or population size in each round. After two rounds, three subsequent sorting rounds were completed by Fluorescent Activated Cell Sorting (FACS) on a Sony SH800 Cell Sorter. Each round, biotinylated FGFR1-D3b and/or biotinylated FGFR1-D3c was used for selection at decreasing concentration. During MACS rounds, Streptavidin Microbeads (Miltenyi Biotec) were coated with biotinylated selection agent prior to mixing with the scFv library. For FACS rounds, the biotinylated selection agent acted as the primary and streptavidin AlexaFluor 488 conjugate (1:500, Life Technologies) as secondary. 1243 Successful scFv display was assessed using chicken anti-cMyc primary (1:500, Life Technologies).

### 4.3 SPR for antibodies

Following MACS and FACS selections, single yeast colonies were plated on SDCAA agar plates, and 96 individual clones were sequenced by Sanger sequencing. Unique clones were reformatted into human IgG1 antibodies using DNA synthesis (Gen-Script). The resulting IgG1 constructs were transiently expressed in CHOK1 cells (30mL cultures) and purified using previously described expression and purification 1257 methods[51–54]. The IgG1 antibodies were evaluated using Surface Plasmon Reso-nance (SPR) on a Biacore T200 (Cytiva) with a Protein A Series S Biosensor chip. The machine was primed with HBS-EP buffer (0.01 M HEPES pH 7.4, 0.15 M NaCl, 3 mM EDTA, 0.005% v/v Surfactant P20), and antigens or antibodies were also diluted into the same buffer for all SPR experiments. Antibodies were captured by Protein A at 2 µg/mL, and Multi-Cycle Kinetics was performed at 25°C to determine binding affinities to both FGFR1-D3b and FGFR1-D3c. All data were analyzed using Biacore Insight software using a 1:1 binding model. A single clone, designated 1C2, was selected for affinity maturation, demonstrating binding affinities of 6.6 nM to FGFR1-D3b and 24.2 nM to FGFR1-D3c.

### 4.4 mRNA display on TEAD

We designed an eight amino acid macrocyclic lariat library in a genetically reprogrammed in vitro translation system as previously described [55–59]. Briefly, we aminoacylated ClAc-L-phenylalanine onto the initiator tRNA^fMet^ and encoded cysteine onto the codon CCA at positions 5, 6, 7 or 8 to spontaneously from athioether link with the cyclic portion of the peptide ranging from a ring size of 4-8. All peptide sequences were followed by a sequence encoding for a G3SGS linker. The remaining positions were randomized with a NNU genetic code containing *N* -methyl-L-phenylalanine (MeF; codon TTC), D-valine (v; codon ATC), tert-Butyl-glycine (tBuG, codon CCT), 4-Phenyl-L-phenylalanine (Bip; codon ACC), (S)-1,2,3,4-Tetrahydroisoquinoline-3-carboxylic acid (Tic; codon GCC), L-4-thiazolyl-Alanine (4TzA, codon GAT), (4-hydroxybenzyl)glycine (HyG, codon TGT) in addition to the eight natural amino acids leucine, valine, serine, glycine, histidine, arginine, asparagine, tyrosine.

Affinity selections of macrocycles binding to TEAD were conducted using biotinylated N-terminal avi-tagged TEAD2-YBD, expressed and purified as described previously [60, 61]. For affinity selections each translation reaction contained 2 *µ*M mRNA-peptide-linker conjugate, 50 *µ*M of each initiator tRNA and 25 *µ*M of each elongator tRNA. Translation reactions were quenched by addition of 17 mM EDTA and reverse transcribed using RNase H minus reverse transcriptase (Promega) at 42°C for 30 minutes. Following a buffer exchange into HBS-T buffer (25 mM HEPES-NaOH pH 7.5, 150 mM NaCl, 0.05% Tween-20), the macrocycle:cDNA library was incubated with 250 nM biotinylated TEAD2-YBD for 20 minutes at room temperature and then incubated with streptavidin-coated beads (Dynabeads M-280 Streptavidin, Thermo Fisher) for 10 minutes. The beads were washed three times with HBS-T. The cDNA was eluted by heating the beads for 5 min at 95°C and the recovery of the elution fraction was determined by qPCR using SYBR Green I on a QuantStudio 5 thermal cycler (Thermo Fisher). The enrichment of peptides for each round was assessed by next-generation sequencing using a HiSeq sequencer (Illumina) of the final cDNA after each round.

### 4.5 SPR for macrocyclic peptides

SPR experiments for macrocyclic peptides were performed by Sygnature Discovery in HBS-T (150 mM NaCl, 50 mM HEPES pH 7.5, 0.001% Tween 20, 0.2% PEG3350) in 2% DMSO. Peptides were diluted six times in a threefold dilution series from 1 mM and injected at 50 mL/min using single-cycle kinetics with 120 s association and 1500 s dissociation. The SPR data was evaluated for binding affinity using the Biacore Insight Software (Cytiva).

### 4.6Evolutionary fitness inference

Evolutionary fitness inference (EVFI) uses input data in the form of a count table **C** with shape *G*×*T*, with *G* unique genotype variants, and *T* total timepoints, and entries in the count table are non-negative integers representing DNA sequencing counts. The main output is an inferred *G*-dimensional fitness vector **w**, where each variant’s fitness is a positive number representing its growth rate over multiple rounds of selection and competition.

Denote mask*_t_*(·) as a function that selects the elements in an input *G*-dimensional vector at indices where *c_g,t_ >* 0. We optimize **w** using maximum likelihood for a likelihood function *ℓ*, which we take to be a Multinomial distribution in this work:

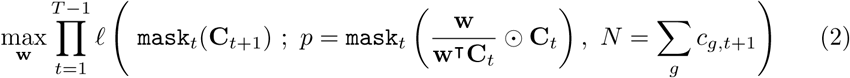

A complete description of EVFI is provided in the Supplementary Material.

### 4.7 DeepEVFI

DeepEVFI jointly learns a sequence-to-fitness deep neural network parametrized by *θ*, and initial abundances of each variant. DeepEVFI first empirically computes a vector ***τ*** where each element is the earliest timepoint where each variant has non-zero count. The predicted abundance of a variant *g* at any time *t* ≥ *τ_g_*is taken to be:

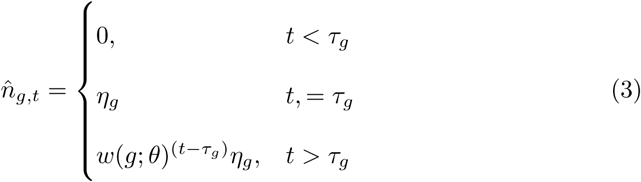

Predicted abundances correspond to predicted frequencies as 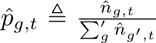. We use a Dirichlet-Multinomial likelihood:

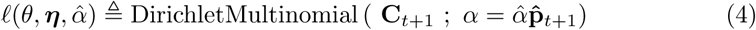

We use a three-stage approach to training DeepEVFI. First, we train EVFI on the input count table to obtain a fitness vector **w**. Second, we warm-up train the deep neural network with mean-squared-error loss on predicting **w**. Third, we perform full end-to-end training, optimizing the neural network weights directly on the count table likelihood loss.

Furthermore, we avoid optimization issues caused by the non-identifiability property by not performing minibatch optimization over variants. However, not all variants may be present in any single timepoint. In practice, we accumulate gradients and take one step per epoch. We describe DeepEVFI methodology in greater detail in the Supplementary Material.

### 4.8 Probability of v-trajectories

We define a *v-trajectory* as a frequency trajectory that decreases, then increases, as in the shape of the letter ‘v’. V-trajectories in true population frequencies are not possible under our assumed model of asexual natural selection. We provide a mathematical proof of this statement in the Supplementary Discussion, and use this property to evaluate how well real-world datasets adhere to our model’s assumptions, which is an important tool in evaluating the trustworthiness of inferred fitness values for decision-making.

Denote a triplet of rounds 1, 2, 3 without loss of generalization and consider variant counts *c*_1_*, c*_2_*, c*_3_, total counts *N*_1_*, N*_2_*, N*_3_ and true population frequencies *p*_1_*, p*_2_*, p*_3_. We compute the probability of a v-trajectory given count data *p*(*p*_1_ *> p*_2_*, p*_2_ *< p*_3_|**c**, **N**) under a binomial noise model *c_t_* ∼ Bin(*p_t_, N_t_*) as:

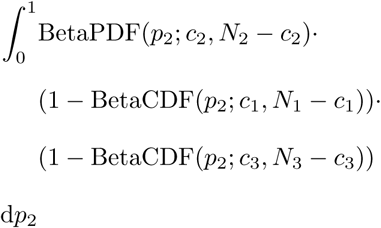

Given count data at more than three rounds, we take the probability a variant has a v-trajectory given all count data, as the maximum probability of a v-trajectory among any triplet of consecutive rounds. To report the fraction of a population comprised of variants violating assumed dynamics, we sum the read counts of all variants with probability of v-trajectory above a threshold, set at 95% in this work.

### 4.9 Method comparison

ACIDES was run with default settings. Enrich2 was run with the ‘full’ argument and WLS scoring for datasets with three or more timepoints, and ratios scoring for datasets with two timepoints. Two-timepoint enrichment was computed for two timepoints *t, t*^′^ for a variant *g* with a pseudocount of 0.5 as:

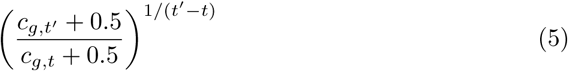

where the power adjustment scales the enrichment between two timepoints which can be separated by several rounds, to the effective enrichment per round.

For evaluation, each methods’ inferred growth rate was used to simulate forward in time all variants present in the second-to-last timepoint population, to the final timepoint, to obtain predicted final timepoint frequencies. This evaluation strategy was chosen as it focuses evaluation on the quality of inferred fitness scores, and deprioritizes the ability to predict which novel variants enter the population, possibly by mutation.

For all methods, the final timepoint was treated as a held-out test set.

To tune hyperparameters for DeepEVFI while using the final timepoint as a heldout test set, we used a two-stage approach. First, we used the 2nd-to-last timepoint as a validation set, and selected hyperparameters based on validation performance on models trained on data before the 2nd-to-last timepoint. Then, we took the best hyperparameters and trained models on data before the final timepoint, and evaluated on the final timepoint. We report DeepEVFI hyperparameters in the Supplementary Material.

Noise-weighted Pearson correlation of predicted enrichment to observed enrichment from time *t* to *t* + 1 is computed with weights for each variant as:

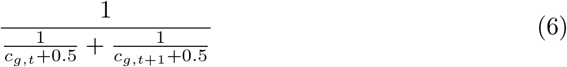

which is motivated as the inverse of the variance of the observed enrichment under Poisson assumptions[16].

### 4.10 Probability of improved fitness

We estimate the probability that variant *g* has higher inferred fitness than variant *r* given count data:

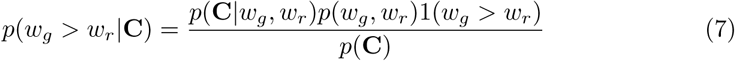

We take *p*(**C**|*w_g_, w_r_*) to follow a Dirichlet-Multinomial distribution, so that:

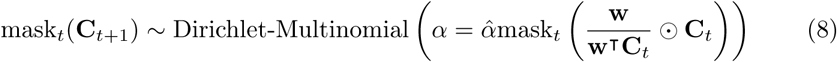

We use a uniform prior for *p*(*w_g_, w_r_*) and learn *α*^ from data by maximum likelihood jointly with fitness inference. To efficiently estimate *p*(*w_g_ > w_r_*|**C**), we use the grouping property of the Dirichlet-Multinomial distribution (Supplementary Note) to group counts into three bins: counts for variant *g*, for variant *r*, and summed counts for all variants other than *g, r* which we denote *o*, whose likelihood are governed by three fitness variables: *w_g_, w_r_, w_o_*. We estimate *p*(*w_g_ > w_r_*|**c_r_**, **c_g_**, **c_o_**) using Monte Carlo estimation. To address the non-identifiability property, we compute likelihoods using uniformly distributed random samples for *w_g_, w_r_* while keeping *w_o_* fixed. We discuss this method in greater detail in Supplementary Note.

### 4.11 Constructing diverse variant sets with high fitness

We use a greedy algorithm that iteratively adds elements to a nomination set. Consider a set of elements *X*, a score function such as log fitness *f* (*x*), and a distance function *d*(*x, x*^′^). The nomination set *S* is initialized with the variant with the highest log fitness, and a weight *α* is chosen. At each iteration, all variants not in the set are scored using:

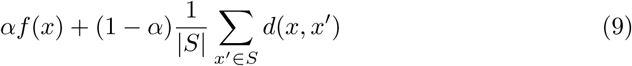

and the highest-scoring variant is added to the set. The iteration repeats until the set size reaches the desired number. To plot the Pareto frontier of nomination sets trading off mean log fitness and mean pairwise distance, we vary *α* from 0 to 1. For all datasets other than M-MP, we use one-hot encoded vectors and L2 distance. For M-MP, we use chirality-aware Morgan fingerprint representations with radius 3 and nBits 2048, and compute distance as 1 -Tanimoto similarity. To make UMAP plots, one-hot encoded vectors were transformed with principal component analysis into up to 50 dimensions, then UMAP was applied.

### 4.12 FGFR1 variant nomination

We used EVFI on three tracks: ABCDE, ABCDF, and ABCDH, using the population names described in the methods section on yeast display on FGFR1. These tracks were chosen as populations E, F, and H underwent selection against both D3b and D3c for the entirety of their selection history. EVFI was used to infer fitness and nominate variants separately for each track, and pooled together. In our study of the correlation between fitness and affinity, we use the fitness scores from the ABCDH track which had the strongest selection pressure.

### 4.13 TEAD variant nomination

To nominate variants, we trained DeepEVFI on the TEAD data, and used to score the peptides. Then, a variational autoencoder was trained on macrocyclic peptides represented with one-hot encoding and Morgan fingerprint using reconstruction loss. The encoder was used to encode the 5000 peptides with highest DeepEVFI score, and UMAP was used to cluster the latent embeddings into 150 clusters. The top-scoring peptide was chosen from each cluster to yield 150 variants, then the top 20 scoring peptides among the 150 were nominated for testing.

