## Supplementary Information for "Deep Evolutionary Fitness Inference for Variant Nomination from Directed Evolution"

**4 Adjusting inferred fitness with off-target data** **10**

**5 Posterior probability of improved fitness** **12**

**6 Mathematical properties of evolutionary fitness inference** **14**

**7 Mutation rates in *in vitro* display assays** **18**

**8 Fitness inference *vs.* computing round-over-round enrichment** **20**

**9 Methods discussion: ACIDES, Enrich2, EVFI** **22**

**10 Experimental details** **24**
10.1 Genotype featurization . . . . . . . . . . . . . . . . . . . . . . . . . . . 24
10.2 DeepEVFI hyperparameters . . . . . . . . . . . . . . . . . . . . . . . . . 24

### 1 Notation

Here, we describe notation used in the methods and supplementary information.

**Table 1** Notation

| | |
|---|---|
| | Variables from Observed Data |
| $G$ | Total number of unique genotype variants in a dataset; a positive integer, indexed by $g$. |
| $T$ | Total number of timepoints in a dataset; a positive integer, indexed by $t$. |
| $c_{g,t}$ | The number of DNA sequencing reads for variant $g$ at timepoint $t$; a non-negative integer. |
| $p_{g,t}$ | The fractional proportion of variant $g$ at timepoint $t$ among all DNA sequencing reads at timepoint $t$. Defined as $c_{g,t}/\sum_{g'} c_{g',t}$. A real number between 0 and 1. |
| | Unobserved Variables |
| $n_{g,t}$ | The absolute quantity of variant $g$ at timepoint $t$, such as number of physical molecules. This is not measurable with DNA sequencing; a non-negative real number. |
| $\omega_g$ | Absolute fitness of variant $g$. A non-negative real number. Not identifiable from data; can only be identified up to an unknown multiplicative factor. |
| | Inference Variables |
| $w_g$ | Inferred fitness, or relative fitness, of variant $g$. A non-negative real number. |
| $\tau_g$ | The timepoint when variant $g$ first enters the population and has non-zero quantity; a positive integer. |
| $\eta_g$ | The absolute quantity of variant $g$ at its initial timepoint $\tau_g$; a non-negative real number. |
| $\phi_g$ | The fractional proportion of variant $g$ at its initial timepoint $\tau_g$. Defined as $\eta_g/\sum_{g'} n_{g',\tau_g}$. A real number between 0 and 1. |
| | Vectors and Matrices |
| $\boldsymbol{\omega}$ | Absolute fitness vector with length $G$, with elements $\omega_g$. |
| $\mathbf{w}$ | Inferred fitness, or relative fitness, vector with length $G$, with elements $w_g$. |
| $\mathbf{p_t}$ | Vector of observed variant frequencies with length $G$, with elements $p_{g,t}$. |
| $\boldsymbol{\tau}$ | Vector of initial timepoints with length $G$, with elements $\tau_g$. |
| $\boldsymbol{\eta}$ | Vector of absolute variant quantities at their initial timepoints, with length $G$, with elements $\tau_g$. |
| $\boldsymbol{\phi}$ | Vector of fractional variant proportions at their initial timepoints, with length $G$, with elements $\tau_g$. |
| $\mathbf{C_t}$ | Vector of time-series DNA sequencing data at timepoint $t$ with length $G$, with elements $c_{g,t}$. |
| $\mathbf{C}$ | Matrix of time-series DNA sequencing data with shape $G \times T$ with elements $c_{g,t}$. |

### 2 Data generative process and problem statement

Here, we state our model of the data generative process and the inference problem.

#### 2.1 Model of data generative process

We consider a process of wet-lab directed evolution, where a population of genotype variants undergo iterated rounds of selection, replication, and potentially mutation. Timepoint samples of the population are taken for DNA sequencing. For $T$ total timepoints and $G$ total unique genotype variants, this process generates a matrix $\mathbf{C}$, which we call a "count table", with entries $c_{g,t}$ denoting the non-zero integer number of DNA sequencing reads for variant $g$ at timepoint $t$ (see Notation section).

We model growth in each round of directed evolution using a simple model of asexual natural selection, where each variant $g$ with absolute quantity $n_{g,t}$ (e.g., absolute abundance of number of molecules, or copies) grows at a rate over time according to its absolute fitness $\omega_g$, a non-negative real number:

$$n_{g,t+1} = \omega_g n_{g,t} \tag{1}$$

We assume that variant $g$ enters the population at most once, at initial timepoint $\tau_g$, with absolute quantity $\eta_g$. Thus, we have:

$$n_{g,t} = \begin{cases} 0, & t < \tau_g \\ \eta_g & t, = \tau_g \\ \omega_g^{(t-\tau_g)} \eta_g, & t > \tau_g \end{cases} \tag{2}$$

We define the population frequency of a variant $g$ at timepoint $t$ as:

$$p_{g,t} \triangleq \frac{n_{g,t}}{\sum_j n_{j,t}} \tag{3}$$

By algebraic manipulation, equation 1 induces non-linear dynamics on variant frequencies (See property 1 for proof):

$$p_{g,t+1} = \frac{\omega_g}{\sum_j \omega_j p_{j,t}} p_{g,t} \tag{4}$$

Finally, we model:

$$\mathbf{C_t} \sim \text{Multinomial}\left(p = \mathbf{p_t}, N = \sum_{g'} c_{g',t}\right) \tag{5}$$

where $c_{g,t}$ is the number of DNA sequencing reads for variant $g$ at timepoint $t$, $\mathbf{C_t}$ is the vectorized version with length $G$, and $\mathbf{p_t}$ is the vector of $p_{g,t}$.

#### 2.2 Inference problem

Given a DNA sequencing count table $\mathbf{C}$ with shape $G \times T$, we aim to infer three $G$-dimensional vectors: the absolute fitness vector $\boldsymbol{\omega}$, and initial timepoints $\boldsymbol{\tau}$ and abundances $\boldsymbol{\eta}$.

However, $\boldsymbol{\omega}$ is not identifiable from measurements of variant frequencies, even if sequencing depth or number of timepoints approach infinity; it is only identifiable up to an unknown multiplicative factor (Property 2). This is because variant frequency dynamics in equation 4 are invariant to rescaling $\omega$ by any positive real number $c$.

As a result, we set up our inference problem as:

**Problem statement.** Given $\mathbf{C}$, infer initial timepoints $\boldsymbol{\tau}$, initial abundances $\boldsymbol{\eta}$, and *relative fitness* $\mathbf{w} \triangleq c\boldsymbol{\omega}$ where $c > 0$ is unknown.

This inference problem is a challenging mixed-integer optimization problem because it includes continuous optimization for $\mathbf{w}, \boldsymbol{\eta}$ and discrete optimization for initial timepoints $\boldsymbol{\tau}$. In principle, as initial timepoints for different genotype variants are independent, $\boldsymbol{\tau}$ can take on up to $T^G$ different values. For typical values $T = 6, G = 30000$, we have $T^G = 2.8 \times 10^{233}$. For this reason, we sought to develop efficient approximate methods.

### 3 EVFI and DeepEVFI

#### 3.1 EVFI: Masked optimization for zero input counts

In EVFI, we focus on inferring relative fitness $\mathbf{w}$ without explicitly inferring $\boldsymbol{\tau}$ and $\boldsymbol{\eta}$ to improve efficiency, while handling their potential impact on estimating $\mathbf{w}$ to retain accuracy.

We observe two properties of the dynamics in equation 4, under the situation where there is no mutation and all variants are present at all timepoints. First, if there is no measurement noise (i.e., $\mathbf{p_t}$ is observed), then $\mathbf{w}$ is identifiable given data from just two timepoints under equation 4. Second, if a variant has zero frequency at time $t$, then that variant will have zero frequency at all following timesteps.

When mutations can introduce variants into populations that follow the dynamics of equation 4, the first property is modified: if there is no measurement noise, then $\mathbf{w}$ is identifiable given data from just two timepoints under equation 4 for all variants that are present in the earlier timepoint. However, variants that are absent, and enter the population at a later timepoint, cannot have their fitness identified.

We exploit these properties to propose a masked optimization procedure, where we iterate over consecutive timepoint pairs $t, t+1$, and optimize our estimate of $\mathbf{w}$ for variants with non-zero count at timepoint $t$. We denote $\texttt{mask}_t(\cdot)$ as a function that selects the elements in an input $G$-dimensional vector at indices where $c_{g,t} > 0$.

We randomly initialize $\mathbf{w}$, and optimize $\mathbf{w}$ using maximum likelihood for a likelihood function $\ell$, which can be a multinomial or dirichlet-multinomial likelihood:

$$\max_{\mathbf{w}} \prod_{t=1}^{T-1} \ell \left( \texttt{mask}_t(\mathbf{C}_{t+1}) \; ; \; p = \texttt{mask}_t(\hat{\mathbf{p}}_{t+1}), \; N = \sum_g c_{g,t+1} \right) \tag{6}$$

where

$$\hat{\mathbf{p}}_{t+1} \triangleq \frac{\mathbf{w}}{\mathbf{w}^\intercal \mathbf{C}_t} \odot \mathbf{C}_t \tag{7}$$

simulates forward in time equation 4 using our current estimate of relative fitness $\mathbf{w}$ and observed variant counts $\mathbf{C}_t$. Note that $\hat{\mathbf{p}}_{t+1}$ is guaranteed to be a probability vector that sums to one, and the equation is equivalent if we replace $\mathbf{C}_t$ with the observed variant frequency vector with elements $c_{g,t} / \sum_g' c_{g',t}$. The symbol $\odot$ indicates element-wise multiplication of two vectors.

We provide python pseudocode for computing the loss over the full dataset in 1 and 2.

Due to the non-identifiability property (Property 2), training is most stable when taking one model update step per epoch on the full-data loss. To do this, we accumulate gradients over minibatches and use it to update the model once the entire dataset has been seen. We use the Adam optimizer to optimize log_fitness until training loss converges (Algorithm 3).

In summary, EVFI sidesteps explicitly modeling when variants enter the population and their initial frequencies to focus on inferring relative fitness, by using masking during optimization to focus on fitness inference using variants that are present in the population.

**Algorithm 1** Loss (full dataset)

```
def fulldata_loss(counts: (T, G), log_fitness: (G) ):
total_loss = 0
for (pre, post) in itertools.pairwise(range(T)) do
    mask = np.where(counts[pre] > 0)
    pred_fqs = predict(mask(counts[pre]), mask(log_fitness), steps=post-pre)
    total_loss += loss(pred_fqs, mask(counts[post]))
end for
return total_loss
```

**Algorithm 2** Predicting variant frequencies

```
def predict(counts: (G), log_fitness: (G), steps: int = 1):
pred_logits = log(counts) + steps * log_fitness
return pred_logits - torch.logsumexp(pred_logits, dim=0)
```

**Algorithm 3** EVFI training

```
def train_evfi(counts: (T, G)):
w: (G) = random_init()
while loss not converged do
    loss = fulldata_loss(counts, w)
    Update w with Adam optimizer using loss
end while
return w
```

#### 3.2 DeepEVFI: Conservative fixed estimate of initial timepoints.

For DeepEVFI, we jointly infer a relative fitness function $w(g;\theta)$ and initial variant abundances $\boldsymbol{\eta}$. We conservatively estimate initial timepoints $\boldsymbol{\tau}$ empirically for each variant as the earliest timepoint where the variant is observed to have non-zero count.

We predict the abundance of a variant $g$ at any time $t \geq \tau_g$ as:

$$\hat{n}_{g,t} = \begin{cases} 0, & t < \tau_g \\ \eta_g & t, = \tau_g \\ w(g;\theta)^{(t-\tau_g)}\eta_g, & t > \tau_g \end{cases} \tag{8}$$

We use predicted abundances to predict frequencies:

$$\hat{p}_{g,t} \triangleq \frac{\hat{n}_{g,t}}{\sum_{g}' \hat{n}_{g',t}} \tag{9}$$

We learn $\theta$ and $\boldsymbol{\eta}$ by maximum likelihood:

$$\max_{\theta,\boldsymbol{\eta},\hat{\alpha}} \prod_{t=1}^{T} \text{DirichletMultinomial}\left(\ \mathbf{C}_{t+1}\ ;\ \alpha = \hat{\alpha}\hat{\mathbf{p}}_{t+1}\right) \tag{10}$$

alongside $\hat{\alpha}$, which is a concentration parameter for the Dirichlet-Multinomial distribution.

We optimize this loss using full-batch gradient descent, as in EVFI, by accumulating gradients over minibatches and updating the model once per epoch. Minibatching over genotype variants as an implementation of minibatch stochastic gradient descent may appear to work here because dynamics equation 4 holds for subsets of genotype variants. However, due to that equation's invariance to rescaling relative fitness (Property 2), each minibatch will learn a different, arbitrary fitness scale which is not directly comparable to the fitness scale of other minibatches. This can substantially hinder training efficiency and the ability to find a good optima in a reasonable amount of time.

**Algorithm 4** DeepEVFI training

```
def train_deepevfi(featurized_genotypes: (G, D), counts: (T, G)):
w: (G) = train_evfi(counts)
model = init_model()

# warmup training
for epoch in num_epochs do
    for minibatch_indices in minibatches do
        pred_w: (G) = model(featurized_genotypes)
        loss = ( pred_w[minibatch_indices] - w[minibatch_indices] )^2
        Update model with Adam optimizer using loss
    end for
end for

# end-to-end training
for epoch in num_epochs do
    pred_w: (G) = model(featurized_genotypes)
    loss = fulldata_loss(counts, pred_w)
    Update model with Adam optimizer using loss
end for
return model
```

In practice, we consider a three-stage approach to train DeepEVFI (Algorithm 4). First, we train standard EVFI on the input count table to obtain a fitness vector $\mathbf{w}$. Second, we warm-up train the deep neural network with mean-squared-error loss on predicting $\mathbf{w}$, using minibatch stochastic gradient descent. Third, we perform full end-to-end training, combining count table likelihood loss with mean-squared-error loss. Denoting the neural network as $f_\theta : \mathcal{X} \rightarrow \mathcal{W}$, the loss in the third stage is:

$$c_0 \ell\left(\theta, \boldsymbol{\eta}, \hat{\alpha}\right) + c_1 \sum_i \left(f_\theta(x_i) - w_i\right)^2 \tag{11}$$

where $c_0, c_1$ are weights that may vary during training. In practice, we used $c_1 = 0$ for our experiments, which discards the regularization term during end-to-end training.

##### 3.3 Remarks.

Here, we remark on our problem framework, the design of EVFI and DeepEVFI, and their properties.

***EVFI supports parallel tracks and directed acyclic graph timepoint relationships.***

This is a property of EVFI, but not DeepEVFI, because EVFI's loss function is decomposable into terms depending only on input/output timepoint pairs ($t, t+1$ conventionally), and EVFI does not explicitly model frequencies unlike DeepEVFI and other methods.

Consider a data structure representation of time series data where each node corresponds to a timepoint, and directed edges connect input timepoints to output timepoints for each round of directed evolution. The simplest type of dataset is where node follow nodes in a single "line" such as $\circ \rightarrow \circ \rightarrow \circ$. However in practice, multiple directed evolution campaigns might be performed on different populations on the same target, yielding parallel tracks: $\circ \rightarrow \circ \rightarrow \circ; \circ \rightarrow \circ \rightarrow \circ$. Alternatively, a population might be split and subjected to two similar new rounds of selection, yielding a bifurcation and a tree structure: $\circ \leftarrow \circ \rightarrow \circ$.

Methods that explicitly model frequencies must learn many sets of initial frequencies on more complex datasets. The total parameter count that is optimized thus can increase with increasingly complex datasets. For instance, parameter count would scale linearly in the number of parallel tracks in a dataset. In contrast, EVFI has a constant number of inference parameters regardless of dataset complexity, because its loss function can be decomposed onto edges in timepoint relationship graph.

***Dirichlet-Multinomial distribution models a form of genetic drift***

The Dirichlet-Multinomial distribution is an overdispersed multinomial distribution, and is equivalent to a hierarchical model where a probability vector $\mathbf{p}$ is drawn from a Dirichlet distribution with parameter $\boldsymbol{\alpha}$, then an observation is drawn from a multinomial distribution with probability vector $\mathbf{p}$ and number of trials $N$.

In the context of evolutionary fitness inference, using a Dirichlet-Multinomial observation distribution can be understood as modeling a form of genetic drift, where there is random variation around each variant's growth from the previous timepoint.

***Empirical properties of directed evolution informing method design***

Although our method seeks to be highly general, our method design is partially motivated by several empirical properties of directed evolution assays such as mRNA display, yeast display, and phage display, which in turn can potentially limit our method's performance on datasets with significantly different properties.

Display assays empirically support very large population sizes of hundreds of millions to trillions of variant units that are acted upon by selection pressure - denote this number $N$. At the same time, mutation rates are relatively low, up to $10^{-5}$ mutations per nucleotide per replication at the highest; denote this $\mu$.

Our simple model of variants entering the population, potentially due to mutation, is motivated by the empirical property that $\mu << 1$, so that in practice, new mutants enter the population at very low frequency, and mutation does not meaningfully reduce the abundance of the parental variant, allowing us to ignore its effect on the parental variant abundance. If the mutation rate was much higher, then explicitly modeling the effect of mutation on the parental variant abundance would likely become necessary.

Large population sizes $N$ can reduce the importance of explicitly modeling genetic drift.

##### *Single-entry assumption is not restrictive*

Our framework assumes that variants enter the population up to one time. With explicit sources of mutation, this assumption can easily be violated in practice; however, we argue here that this violation is not restrictive and makes little difference for fitness inference.

In property 4, we prove that in our framework, frequency-weighted fitness increases monotonically over time. In property 5, we prove in our framework, v-trajectories are not possible, which is when a variant decreases in frequency then increases in frequency.

Taken together, we can see that even if a variant truly does enter the population a second time, if the population follows the assumed dynamics, then the variant has low fitness, because it previously entered the population, but was outcompeted. When the variant enters the population a second time, the population has only become more competitive (its frequency-weighted fitness is now higher), so the variant will continue to be outcompeted. While accurate fitness inference for such a variant can be challenging when its frequencies are so low, our methods will generally infer low fitness for such variants.

##### *Accounting for time-varying selection pressure*

In our framework, we assume that fitness does not vary by time. A simple way to account for time-varying selection pressure in this framework is to artificially add more "time" or "generations" between timepoints that have relatively higher selection pressure.

### 4 Adjusting inferred fitness with off-target data

A common setting is availability of directed evolution data for a single population evolved in (a) a campaign against the target of interest and the instrument (target + instrument), and also (b) a campaign only against the instrument (instrument only). For example, a target protein may be immobilized on beads (representing the instrument), and the evolved drugs are washed over the bead-bound immobilized target protein, during the (target + instrument) campaign. Separately, the population can be split and washed only over the beads, without the target protein present, to attempt to adjust for general stickiness. With such data, two enrichment scores or fitnesses can be estimated; a (target + instrument) fitness, and an (instrument-only) fitness.

These values are commonly combined by dividing the (target + instrument) fitness by the (instrument-only) fitness: genotype variants with a high ratio score thus have relatively higher (target + instrument) fitness, and relatively lower (instrument-only) fitness, making them good candidates for variant nomination and further testing.

However, this approach fails to consider the identifiability up to proportionality property of fitness. As two fitness inference procedures were performed, the relative scale between the two inferred fitnesses is unrelated and unknown. For example, it may be that no genotype variant has any meaningful off-target activity, or that all genotype variants do. These two scenarios would lead to dramatically different decisions in practice: perhaps the entire population is no good, or perhaps we can mine deeper into the population than otherwise expected. These scenarios can be distinguished by measuring the total absolute abundance of the population: if it decreased dramatically when washing only over beads, then the population as a whole has little off-target activity. However, DNA sequencing data of population frequencies does not provide such information. Prior work and common practice fails to appreciate this rescaling issue, which may lead to suboptimal decision-making.

Here, we prove an upper-bound as a partial solution to this rescaling problem under the assumption that (target + instrument) fitness is a sum of (target-only) fitness and (instrument-only) fitness. This upper bound on the rescaling factor provides a simple, principled method to correct fitnesses for off-target activity, and distinguish between populations with high or low aggregate off-target activity, enabling scientists to mine deeper or choose to discard populations.

**Theorem 1** (Upper bound for adjusting off-target fitness). *Suppose, from inferred relative fitness* $c_1\mathbf{w}_{total}$ *and* $c_2\mathbf{w}_{off}$ *which are proportional by unknown constants* $c_1, c_2$ *to absolute fitness* $\boldsymbol{\omega}$ *as* $c_1\mathbf{w}_{total} = \boldsymbol{\omega}_{total}$ *and* $c_2\mathbf{w}_{off} = \boldsymbol{\omega}_{off}$*. Further, suppose that*

$$\boldsymbol{\omega}_{total} = \boldsymbol{\omega}_{on} + \boldsymbol{\omega}_{off}. \tag{12}$$

*Rewritten, we have:*

$$\underbrace{c_1\mathbf{w}_{total}}_{available} = \underbrace{c_1\mathbf{w}_{on}}_{goal} + \underbrace{\frac{c_1}{c_2}}_{unknown} \underbrace{c_2\mathbf{w}_{off}}_{available}. \tag{13}$$

*Then,*

$$\arg\min_{g} \frac{c_1 w_{total,g}}{c_2 w_{off,g}} \geq \frac{c_1}{c_2}. \tag{14}$$

*Proof.* Using the property that absolute fitness must be non-negative,

$$c_1 \mathbf{w}_{\texttt{total}} = c_1 \mathbf{w}_{\texttt{on}} + \frac{c_1}{c_2} c_2 \mathbf{w}_{\texttt{off}}$$
$$c_1 \mathbf{w}_{\texttt{total}} \geq \mathbf{0} + \frac{c_1}{c_2} c_2 \mathbf{w}_{\texttt{off}}$$

Dividing through, for each element $g$ we have:

$$\frac{c_1 w_{\texttt{total},i}}{c_2 w_{\texttt{off},i}} \geq \frac{c_1}{c_2}.$$

□

#### 5 Posterior probability of improved fitness

In this section, we discuss our approach for estimating the probability that variant $g$ has higher fitness than a reference variant $r$ given count table data, which is expressed as:

$$p(w_g > w_r|\mathbf{C}) = \frac{p(\mathbf{C}|w_g, w_r)p(w_g, w_r)1(w_g > w_r)}{p(\mathbf{C})} \tag{15}$$

where $1(\cdot)$ is the indicator function.

We take $p(\mathbf{C}|w_g, w_r)$ to follow a Dirichlet-Multinomial distribution, so that:

$$\text{mask}_t(\mathbf{C}_{t+1}) \sim \text{Dirichlet-Multinomial}\left(\alpha = \hat{\alpha}\text{mask}_t(\hat{\mathbf{p}}_{t+1})\right) \tag{16}$$

where

$$\hat{\mathbf{p}}_{t+1} \triangleq \frac{\mathbf{w}}{\mathbf{w}^\intercal \mathbf{C}_t} \odot \mathbf{C}_t \tag{17}$$

We use a uniform prior for $p(w_g, w_r)$ and learn $\hat{\alpha}$ from data by maximum likelihood jointly with fitness inference.

Computing $p(\mathbf{C}|w_g, w_r)$ which marginalizes over the fitness values of all variants besides $g, r$ is a challenging problem involving very high-dimensional integration in the typical case where the number of unique genotype variants is 30,000. However, the aggregation property of the Dirichlet-Multinomial distribution holds that if $(x_1, ..., x_k) \sim \text{DM}(\alpha_1, ..., \alpha_k)$, then for any indices $i, j$, $(x_1, ..., x_i + x_j, ..., x_k) \sim \text{DM}(\alpha_1, ..., \alpha_i + \alpha_j, ..., \alpha_k)$. We use this property to construct a likelihood for $w_g, w_r$ that is easier to compute by aggregating variants into three bins: $g$, $r$, and all other variants which we denote $o$ for "other". Aggregation is performed by summation, so that

$$c_{o,t} \triangleq \sum_{i \notin \{g,r\}} c_{i,t}. \tag{18}$$

Dropping the mask notation for ease of exposition, and applying aggregation, yields:

$$p(c_{g,t}, c_{r,t}, c_{o,t}|w_g, w_r, w_o) = \text{Dirichlet-Multinomial}\left(\alpha = \hat{\alpha}(\hat{p}_{g,t}, \hat{p}_{r,t}, \hat{p}_{o,t})\right) \tag{19}$$

As predicted frequencies $\hat{p}$ are a function of count data in the previous timepoint, the aggregated likelihood is a product over all timepoints:

$$p(\mathbf{c_g}, \mathbf{c_r}, \mathbf{c_o}|w_g, w_r, w_o) = \prod_{t=2}^{T} p(c_{g,t}, c_{r,t}, c_{o,t}|w_g, w_r, w_o) \tag{20}$$

As this likelihood is invariant to scaling $w_g, w_r, w_o$ together by any multiplicative constant (Property 2), we hold $w_o$ fixed. We use $N$ uniformly distributed random samples of $w_g, w_r$ to estimate:

$$p(w_g > w_r|\mathbf{C}) \approx \frac{\sum_{i=1}^{N} p(\mathbf{c_g}, \mathbf{c_r}, \mathbf{c_o}|w_{g,i}, w_{r,i}, w_o) 1(w_{g,i} > w_{r,i})}{\sum_{i=1}^{N} p(\mathbf{c_g}, \mathbf{c_r}, \mathbf{c_o}|w_{g,i}, w_{r,i}, w_o)} \tag{21}$$

We handle masking in our computational implementation of this approach.

### 6 Mathematical properties of evolutionary fitness inference

In this section, we present mathematical properties of our data generative process, and evolutionary fitness inference.

We begin by restating our simple model of growth under asexual natural selection from equation 1.

$$n_{g,t+1} = \omega_g n_{g,t} \tag{22}$$

---

**Property 1** (Non-linear population frequency dynamics)**.** *Suppose a population follows the dynamics of equation 1. Then, population frequency* $p_{g,t} \triangleq n_{g,t} / \sum_g' n_{g',t}$ *obeys:*

$$p_{g,t+1} = \frac{w_g}{\sum_j w_j p_{j,t}} p_{g,t} \tag{23}$$

*or equivalently, written in bold vector notation,*

$$\mathbf{p_{t+1}} = \frac{\mathbf{w}}{\mathbf{w}^\intercal \mathbf{p_t}} \odot \mathbf{p_t} \tag{24}$$

*where* $\odot$ *means element-wise multiplication.*

*Proof.* Using $N_{t+1} = \sum_g n_{g,t+1} = \sum_g w_g n_{g,t}$, we have

$$\begin{aligned}
n_{g,t+1} &= w_g n_{g,t} \\
\frac{n_{g,t+1}}{N_{t+1}} &= \frac{w_g}{N_{t+1}} n_{g,t} && \text{(divide by } N_{t+1}) \\
p_{g,t+1} &= \frac{w_g}{\sum_j w_j n_{j,t}} n_{g,t} && \text{(replace with definitions)} \\
p_{g,t+1} &= \frac{N_t w_g}{\sum_j w_j n_{j,t}} \frac{n_{g,t}}{N_t} && \text{(multiply by } N_t/N_t) \\
p_{g,t+1} &= \frac{w_g}{\sum_j w_j \frac{n_{t,j}}{N_t}} p_{g,t} && \text{(move } N_t \text{ through fraction)} \\
p_{g,t+1} &= \frac{w_g}{\sum_j w_j p_{j,t}} p_{g,t} && \text{(replace with definitions)}
\end{aligned}$$

□

**Remark.** In vector notation, $\mathbf{w}, \mathbf{p_t}, \mathbf{p_{t+1}}$ are all $G$-dimensional vectors where $\mathbf{p_t}, \mathbf{p_{t+1}}$ have positive entries summing to 1, and $\mathbf{w}$ has positive entries.

Thus far we have considered discrete time dynamics. The continuous time analogue, for time update $\Delta t$, is:

$$\mathbf{p_{t+\Delta t}} = \frac{\exp(\Delta t \log(\mathbf{w}))}{\exp(\Delta t \log(\mathbf{w}))^\intercal \mathbf{p_t}} \odot \mathbf{p_t}. \tag{25}$$

---

**Property 2** (Fitness scale invariance, or identifiability up to proportionality)**.** *In equation 24, multiplying* $\mathbf{w}$ *by a positive constant factor* $c$ *for a given* $\mathbf{p_t}$ *does not change* $\mathbf{p_{t+1}}$*.*

*Proof.*

$$\begin{aligned}\mathbf{p_{t+1}} &= \frac{c\mathbf{w}}{(c\mathbf{w})^\intercal \mathbf{p_t}} \odot \mathbf{p_t} \\ &= \frac{c\mathbf{w}}{c\mathbf{w}^\intercal \mathbf{p_t}} \odot \mathbf{p_t} \\ &= \frac{\mathbf{w}}{\mathbf{w}^\intercal \mathbf{p_t}} \odot \mathbf{p_t}\end{aligned}$$

□

**Remark.** Property 2 means that given a count table, the scale of fitness values cannot be determined, so only the relative ratio between any two fitness values is meaningful. If fitness values are not explicitly scaled during training, then it should be expected that running fitness inference multiple times on the same dataset will give differently scaled fitness values. This is an essential property of fitness inference with significant impact on interpreting and using fitness values for downstream applications.

---

**Property 3** (Population dynamics also hold for population subsets)**.** *Suppose* $\mathbf{w}, \mathbf{p_t}, \mathbf{p_{t+1}}$ *are* $G$*-dimensional vectors for* $G$ *genotype variants that obey equation 24. For any* $m$*-dimensional subset of these genotype variants with associated subset vectors* $\mathbf{w}_s, \mathbf{p}_{s,t}, \mathbf{p}_{s,t+1}$ *(where* $\mathbf{p}_{s,t}, \mathbf{p}_{s,t+1}$ *do not sum to one, as they are subsets of a frequency vector), denote* $\tilde{\mathbf{p}}_{s,t} \triangleq \mathbf{p}_{s,t}/(\sum_{j=1}^{m} p_{s,t,j})$ *as the re-normalized* $m$*-dimensional vector of frequencies of the subset. Then, we have:*

$$\tilde{\mathbf{p}}_{s,t+1} = \frac{\mathbf{w}_s}{\mathbf{w}_s^\intercal \tilde{\mathbf{p}}_{s,t}} \odot \tilde{\mathbf{p}}_{s,t} \tag{26}$$

*Proof.* Without loss of generality, suppose $\mathbf{w}, \mathbf{p_t}, \mathbf{p_{t+1}}$ are ordered such that the subset $s$ corresponds to the first $m$ elements of each. We have for each $i$ where $1 \leq i \leq m$:

$$\begin{aligned}p_{t+1,i} &= \frac{w_i}{\sum_{j=1}^{G} w_j p_{t,j}} p_{t,i} \\ \frac{p_{t+1,i}}{\sum_{k=1}^{m} p_{t+1,k}} &= \frac{w_i}{\sum_{j=1}^{G} w_j p_{t,j}} p_{t,i} \left(\frac{1}{\sum_{k=1}^{m} p_{t+1,k}}\right) \\ \frac{p_{t+1,i}}{\sum_{k=1}^{m} p_{t+1,k}} &= \frac{w_i}{\cancel{\sum_{j=1}^{G} w_j p_{t,j}}} p_{t,i} \left(\frac{\cancel{\sum_{j=1}^{G} w_j p_{t,j}}}{\sum_{k=1}^{m} w_k p_{t,k}}\right)\end{aligned}$$

where we used $\sum_{k=1}^{m} p_{t+1,k} = \sum_{k=1}^{m} \frac{w_k}{\sum_{j=1}^{G} w_j p_{t,j}} p_{t,k}$. Cancelling the term $\sum_{j=1}^{G} w_j p_{t,j}$, we obtain

$$\frac{p_{t+1,i}}{\sum_{k=1}^{m} p_{t+1,k}} = \frac{w_i}{\sum_{k=1}^{m} w_k p_{t,k}} p_{t,i}$$
$$= \frac{w_i}{\sum_{k=1}^{m} w_k \frac{p_{t,k}}{\sum_{z=1}^{m} p_{t,z}}} \frac{p_{t,i}}{\sum_{k=1}^{m} p_{t,k}}$$
$$\tilde{p}_{t+1,i} = \frac{w_i}{\sum_{k=1}^{m} w_k \tilde{p}_{t,k}} \tilde{p}_{t,i}$$

□

---

**Property 4** (Frequency-weighted fitness increases monotonically over time)**.** *Suppose* $\mathbf{w}$ *and* $\mathbf{p_t}$ *for rounds* $t$ *are* $G$*-dimensional vectors for* $G$ *genotype variants that obey equation 24. For any* $t$*, we have:*

$$\mathbf{w}^\intercal \mathbf{p_{t+1}} \geq \mathbf{w}^\intercal \mathbf{p_t} \tag{27}$$

*Proof.* By equation 24, we have:

$$p_{g,t+1} = \frac{w_g}{\sum_j w_j p_{j,t}} p_{g,t} \tag{28}$$

Intuitively, consider three cases: for any variant indexed $g$:

1. $w_g > \mathbf{w}^\intercal \mathbf{p_t}$: Then $\frac{w_g}{\sum_j w_j p_{j,t}} > 1$, and $p_{g,t+1} > p_{g,t}$.
2. $w_g = \mathbf{w}^\intercal \mathbf{p_t}$: Then $\frac{w_g}{\sum_j w_j p_{j,t}} = 1$, and $p_{g,t+1} = p_{g,t}$.
3. $w_g < \mathbf{w}^\intercal \mathbf{p_t}$: Then $\frac{w_g}{\sum_j w_j p_{j,t}} < 1$, and $p_{g,t+1} < p_{g,t}$.

Because $\mathbf{p_{t+1}}$ and $\mathbf{p_t}$ both sum to 1, the net effect is $\mathbf{p_{t+1}}$ places higher weight on all $w_g$ greater than $\mathbf{w}^\intercal \mathbf{p_t}$, relative to $\mathbf{p_t}$. It places lower weight on all $w_g$ less than $\mathbf{w}^\intercal \mathbf{p_t}$. And it places the same weight ($p_{g,t+1} = p_{g,t}$) on all $w_g = \mathbf{w}^\intercal \mathbf{p_t}$. Thus, we can intuitively see that $\mathbf{w}^\intercal \mathbf{p_{t+1}} \geq \mathbf{w}^\intercal \mathbf{p_t}$.

More formally, we rewrite the lemma, expressing $\mathbf{p_{t+1}}$ in terms of $\mathbf{p_t}$:

$$\sum_g w_g \frac{w_g p_{g,t}}{\sum_j w_j p_{j,t}} \geq \sum_g w_g p_{g,t} \tag{29}$$

For clarity of notation, we drop $t$ moving forward:

$$\sum_i w_i^2 p_i \geq \left( \sum_i w_i p_i \right)^2$$
$$\sum_i w_i^2 p_i \geq \sum_i w_i^2 p_i^2 + \sum_j^G \sum_{k>j}^G 2 w_j w_k p_j p_k$$

$$\sum_i w_i^2 p_i(1-p_i) \geq \sum_j^G \sum_{k>j}^G 2w_j w_k p_j p_k$$

$$\sum_i \left( w_i^2 p_i \left( \sum_{j \neq i} p_j \right) \right) \geq \sum_j^G \sum_{k>j}^G 2w_j w_k p_j p_k$$

$$\sum_j \sum_{k>j} w_j^2 p_j p_k + w_k^2 p_j p_k \geq \sum_j^G \sum_{k>j}^G 2w_j w_k p_j p_k$$

$$\sum_j \sum_{k>j} w_j^2 p_j p_k - 2w_j w_k p_j p_k + w_k^2 p_j p_k \geq 0$$

$$\sum_j \sum_{k>j} p_j p_k (w_j - w_k)^2 \geq 0$$

which is true, as $p_i > 0$ for all $i$ by definition.

□

---

**Property 5** (V-trajectories are not possible: variant frequencies cannot decrease then increase)**.** *Suppose* $\mathbf{w}$ *and* $\mathbf{p_t}$ *for rounds* $t$ *are* $G$*-dimensional vectors for* $G$ *genotype variants that obey equation 24. Then, for any variant* $g$ *and consecutive time rounds* $t < t' < t''$*, it cannot be true that* $p_{g,t} > p_{g,t'}$ *and* $p_{g,t'} < p_{g,t''}$*.*

*Proof.* We use the property that frequency-weighted fitness $\mathbf{w}^\intercal \mathbf{p_t}$ increases monotonically over time (equation 27). As

$$p_{g,t+1} = \frac{w_g}{\sum_j w_j p_{j,t}} p_{g,t} \tag{30}$$

the rate of change in frequency $\frac{w_g}{\sum_j w_j p_{j,t}}$ decreases monotonically over time, making the frequency trajectory a concave function over time. □

**Remark.** This theorem states that under a simple model of exponential growth, variant frequencies cannot decrease then increase (a ”v-trajectory”). However, real-world datasets can have v-trajectories for a variety of reasons, such as changes in selection pressure (i.e., changing targets). If a population undergoes extreme bottlenecking between some rounds, then v-trajectories can occur due to high-variance genetic drift. It is possible for the frequency-weighted fitness of a population to decrease if many low-fitness variants are introduced into a population, which can induce v-trajectories when analyzing the population as whole. This particular issue can be avoided during data analysis by instead searching for v-trajectories for a query variant within a restricted subset of variants that all first appear in the population at the same time or earlier than the first appearance of the query variant.

### 7 Mutation rates in *in vitro* display assays

In this section, we discuss the possible rates of mutation that may occur in three common *in vitro* display protocols: yeast surface display, phage display, and mRNA display. We express mutation rates in units of mutations per nucleotide per replication, which we abbreviate as mut/nt/rep.

##### *Phage display*

A common protocol for phage display uses M13 filamentous phage replicating inside *E. coli*. M13 uses the host *E. coli* DNA polymerase to replicate its phage genome. *E. coli* DNA polymerase replicating the *E. coli* genome has an overall mutation rate of $10^{-10}$ mut/nt/rep. However, M13 replicating using *E. coli* DNA polymerase has been observed by multiple groups to have 1000× higher mutation rate, at around $10^{-7}$ mut/nt/rep [1–3]. This fits Drake's rule, at 0.003 muts/genome/rep [3].

In general, DNA replication is a complex process involving DNA polymerase but also multiple DNA repair pathways involving a variety of proteins. *E. coli*'s replication mutation rate of $10^{-10}$ mut/nt/rep can be attributed to three mechanisms: base selection with $10^5$-fold reduction in mutation rate, mismatch repair with $10^3$-fold reduction, and proofreading with $10^2$-fold reduction [4]. Thus, *E. coli*'s high-fidelity replication is attributable not just to the use of *E. coli* DNA polymerase but many other DNA repair proteins.

In principle, it is not clear that M13 phage replicating using *E. coli* DNA polymerase would also benefit from exactly the same DNA repair machinery. Previous work has observed that the phage genome is depleted for GATC motifs which contribute to methylation-induced mismatch repair in *E. coli*, which may explain increased observed mutation rates in phage [5].

Furthermore, phage generation time is fast, with M13 modeled at 30 minutes per generation. In practice, many generations of phage replication might occur between each selection round in phage display, which increases the effective mutation rate. For instance, if phage are left to replicate in *E. coli* overnight (12 hours) such that M13 phage replicate 24 times, the rate of $10^{-7}$ mut/nt/rep becomes $2.4 \times 10^{-6}$ mut/nt/24-reps.

Extending this to an evolving gene variant with 100 nucleotides and a library size of $10^9$ molecules, about $10^5$ mutations are introduced across the population every 12 hours of phage replication on average.

##### *mRNA display*

In mRNA display, polymerase chain reaction (PCR) can be used between selection rounds to amplify the population. The ThermoFisher Phusion High-Fidelity DNA polymerase has a mutation rate of $4.4 \times 10^{-7}$ mut/nt/rep. While the number of PCR cycles used in practice can vary, over 30 replication cycles, this becomes $1.3 \times 10^{-5}$ muts/nt. If the evolving peptide or gene has a length of 100 nucleotides, and the population has $10^{13}$ molecules in it, a mutation rate of $1.3 \times 10^{-5}$ muts/nt means $10^{10}$ (ten billion) mutations are introduced across the population each round, on average.

Not using a high-fidelity DNA polymerase can raise mutation rates significantly. *Taq* DNA polymerase has a mutation rate of $2.28 \times 10^{-5}$ mut/nt/rep, which is nearly 100 times higher than the ThermoFisher Phusion High-Fidelity DNA Polymerase.

##### *Yeast display*

A common protocol for yeast surface display uses pCTcon2 plasmid in *Saccharomyces cerevisiae*. The pCTcon2 plasmid is a low copy number plasmid typically with one copy per yeast cell, which helps to ensure an accurate connection between the phenotype that selection acts upon, and the genotype that induces the phenotype. pCTcon2 is a centromeric plasmid which replicates like an additional yeast chromosome. Diploid *S. cerevisiae* genome replication occurs at $10^{-10}$ mut/nt/rep [6], while haploid genome replication has been reported at $10^{-9}$ mut/nt/rep [7]. pCTcon2 functioning as a single-copy chromosome is expected to inherit the haploid mutation rate.

Yeast doubling time is about 90 minutes. If yeast are grown for 24 hours which is 16 generations, the $10^{-9}$ mut/nt/rep becomes $1.6 \times 10^{-8}$ mut/nt/24hr.

With an evolving gene length of 100 nucleotides and a library size of $10^{8}$, a mutation rate of $1.6 \times 10^{-8}$ mut/nt/24hr means that 100 mutations are introduced across the population each round, on average.

##### *Additional sources of mutation*

Physical stress can cause mutations. Furthermore, there can be significant variation in mutation rates: hot spots and cold spots in the genome can vary by up to $10\times$ [8]. Environment and growth conditions can also impact mutation rates, by up to 3-fold [9].

### 8 Fitness inference *vs.* computing round-over-round enrichment

In our framework, if the data comprises two timepoints, fitness inference is conceptually identical to computing round-over-round enrichment.

Round-over-round enrichment for a variant $g$ is computed as:

$$\text{enrichment}(g) = \frac{p_{g,t=1}}{p_{g,t=0}}. \tag{31}$$

so the enrichment ratio comparing two variants $g, r$ is:

$$\frac{\text{enrichment}(g)}{\text{enrichment}(r)} = \frac{p_{g,t=1}}{p_{g,t=0}} \frac{p_{r,t=0}}{p_{r,t=1}} \tag{32}$$

In our framework, fitness dynamics on variant frequencies follow

$$p_{g,t+1} = \frac{\omega_g}{\sum_j \omega_j p_{j,t}} p_{g,t} \tag{33}$$

so the fitness ratio comparing two variants $g, r$ is:

$$\frac{\omega_g}{\omega_r} = \frac{p_{g,t=1}}{p_{g,t=0}} \frac{p_{r,t=0}}{p_{r,t=1}} \tag{34}$$

This relationship is true for any choice of two variants, so it holds for all variants.

Here, we discuss the differences between fitness inference and round-over-round enrichment.

##### *Fitness is a single framework for using all timepoint data, simplifying decision-making*

The primary difference is that fitness inference provides a framework and method that supports three or more timepoints, while round-over-round enrichment requires prespecifying two timepoints. When $T$ timepoints are available, there are $\binom{n}{k}$ ordered timepoint pairs that can be used to compute different round-over-round enrichments. For a typical $T = 5$, there are 10 timepoint pairs: $1 \rightarrow 5, 2 \rightarrow 5, ..., 4 \rightarrow 5$. In general, enrichment computed using different timepoint pairs will disagree with each other – some variants will score highly using certain timepoints pairs, but not others – making it unclear how to make decisions to nominate variants.

##### *Fitness denoises round-over-round enrichment*

The final timepoint represents the most competitive selection environment, so in the above example, $4 \rightarrow 5$ would be the best timepoint pair to compute enrichment to identify variants with highest fitness. However, by using all available timepoints, fitness inference can denoise enrichment scores. For example, two variants might have an enrichment ratio of 2 in the final timepoint pair, but enrichment ratios of 1.5 and 1.7 in earlier rounds. Only using round-over-round enrichment would only use one of these values, while fitness inference would effectively average over these observed enrichment ratios to estimate a denoised version of enrichment.

##### ***Round-over-round enrichment is confounded by zeros, while fitness inference gracefully handles zeros***

With round-over-round enrichment, it is unclear what to do when the input read count for a variant is zero, as it is not possible to divide by zero. Some methods add pseudocounts to treat zeros as very small numbers, but this can introduce a significant source of bias in follow-up decision-making with little supporting evidence.

##### ***Fitness enables comparing variants that do not directly compete with each other, and scores more variants***

EVFI is capable of inferring fitness for any variant as long as it has non-zero counts in at least one pair of consecutive timepoints. Furthermore, EVFI infers fitness for all such variants on the same scale.

Suppose a variant has read count data over 5 timepoints: $[0, 0, 0, 3, 10]$, while another variant has $[5, 3, 0, 0, 0]$. Under very mild conditions – there is a "bridge" of variants that compete with each other, that connects the first two timepoints to the last timepoints – fitness inference will infer fitness of both variants and enable comparing these two variants on the same scale.

In contrast, round-over-round enrichment will only be able to compute a meaningful, data-driven enrichment score for each variant in one out of ten possible timepoint pairs, and cannot compute enrichment scores to compare the two variants.

Because of EVFI's ability to handle zeros in this manner, the size of the variant set with inferred fitness will always be equal to or greater than the size of the variant set with data-driven enrichment scores for any timepoint pair.

### 9 Methods discussion: ACIDES, Enrich2, EVFI

Here, we discuss differences between ACIDES, Enrich2, and EVFI.

***Enrich2 does not properly account for non-linear effects at three or more timepoints***

Enrich2 is primarily designed for deep mutational scanning which compares enrichment of mutants around a wild-type sequence. Its method differs based on whether there is a designated wild-type sequence or not, and Enrich2 generally assumes that the wild-type sequence is observed in every timepoint. In general, directed evolution acts on a diverse input population that has no clearly designated wild-type sequence, so we study the procedure that Enrich2 uses for datasets with three or more timepoints lacking a wild-type sequence. In our notation, Enrich2 performs regression for a variant $g$ on a paired dataset $(t, y(t))^T$, where the regression target $y(t)$ is:

$$y(t) \triangleq \log\left(\frac{\frac{1}{2} + c_{g,t}}{\frac{1}{2} + \sum_{g'} c_{g',t}}\right) \tag{35}$$

$$\approx \log(p_{g,t}) \tag{36}$$

where $\frac{1}{2}$ acts as a pseudocount, and in the second line removing the pseudocount we see that Enrich2 effectively regresses on log variant frequency, taking the regression slope as the variant's score.

Enrich2 assumes that variant score does not vary by time, which means it assumes the same dynamics in equation 4 which we use in EVFI. However, Enrich2's method for score regression contradicts this premise: in the noiseless setting, we show its regression score is not consistent with $\boldsymbol{\omega}$ in equation 4.

Taking the log of equation 4, we get:

$$\log p_{g,t+1} - \log p_{g,t} = \log w_g - \log \sum_{g'} w_{g'} p_{g',t} \tag{37}$$

Enrich2's regression slope effectively captures the right hand side, which includes the term $\log \sum_{g'} w_{g'} p_{g',t}$ which is a function of $t$. For the regression slope to correspond to log fitness, it would have to be equivalent to $\log w_g$, but it is not.

This shows that despite Enrich2 assuming the dynamics in equation 4, and despite aiming to infer a single enrichment score corresponding to fitness, its method is not correct in inferring a score that corresponds to fitness.

***ACIDES has poor behavior on variants with many zero counts***

Consider a variant which has zero read count in every timepoint except the last timepoint, where it has a read count of one. ACIDES assumes all variants are present at all timepoints, and infers an initial abundance $\rho_{g,t=1}$ for each variant $g$ at the first timepoint, and a fitness score $a_g$ representing the rate of growth under selection pressure.

ACIDES models the expected frequency of variant $g$ at time $t$ as:

$$\rho_{g,t} = \frac{\rho_{g,t=1}\exp(a_g t)}{\sum'_g \rho_{g',t=1}\exp(a_{g'} t)} \tag{38}$$

which we recognize as the continuous-time version of equation 4 when $a_g$ corresponds to log fitness. ACIDES learns $a_g, \rho_{g,t=1}$ from count data by maximum likelihood.

For a variant which has zero read count in all timepoints except the last timepoint, the maximum likelihood solution will push initial frequency as close to zero as possible, and push the growth rate $a_g$ to be as high as possible. This pushes the expected frequencies as close to zero as possible for all timepoints other than the last timepoint. This can cause ACIDES to infer excessively high scores for these variants which have little observed data.

Importantly, we observe variants like these in multiple datasets we study in this work. Such variants are therefore not uncommon.

ACIDES' rank robustness metric can assist in decision-making for variant prioritization and serves to add large error bars and uncertainty around the score estimates for such variants. However, error estimates and confidence intervals are constructed using a Gaussian approximation to the likelihood function centered at the maximum likelihood estimate. This means that even though score estimates can have high uncertainty, it is symmetric around artificially high score estimates.

### 10 Experimental details

#### 10.1 Genotype featurization

Datasets C, D, F, Y-Ab, and M-MP had genotype information. Here, we describe how we featurized these genotypes for input to DeepEVFI.

- Dataset C: Phage display, antibody IgH. 12-mer DNA. Examples genotypes: TTG-GTTCCGGAT, TTTGGTTGTCTG, CAGGGGCTGGGG. Featurization: 12*5 = 60-dimensional one-hot encoding.
- Dataset D: Phage display, hYAP65 WW. 34-mer amino acids. Example genotypes: DVPLPAGWEMAKTSSGQRYFLNHIDQTTTWQDPR, DVPLPAGWEMAKTSSGQRYFLNHIDQTTTRQDPR, DVPLPAGWEMAKTSS-GQRYFLNHIDHTTTWQDPR. Featurization: 34*21 = 714-dimensional one-hot encoding.
- Dataset F: Yeast growth, U3 snoRNA. Approximately 20-mer DNA, with variable-length up to 29-mers. Example genotypes: TACTCCGACCCGCCGTATT, CTAACCTTTCCCCATTGTCC, TTTAAGTCATCCAAATCCTC. Featurization: Padded up to 29-mers. 29*5 = 145-dimensional one-hot encoding.
- Dataset Y-Ab: Yeast display, antibodies. Approximately 55-mer amino acids. Featurization: 55*21 = 1155-dimensional one-hot encoding.
- Dataset M-MP: mRNA display, macrocyclic peptides. 8-mer amino acids. Featurization: Converted SMILES to Morgan fingerprints: 128-bit, chirality-aware.

#### 10.2 DeepEVFI hyperparameters

We report the hyperparameters for DeepEVFI models in table 2. Each dataset varied significantly in input dimension size, input data type, number of genotypes, number of timepoints, noise in the dataset, and complexity of the sequence-fitness relationship. As such, a reasonable expectation *a priori* is that the best model size and hyperparameters may also vary significantly, but surprisingly, this was not the case.

As discussed in the methods, we tuned hyperparameters while keeping the final timepoint as a held-out test set by using a two stage approach. First, we used the 2nd-to-last timepoint as a validation set, and selected hyperparameters based on validation performance on models trained on data before the 2nd-to-last timepoint. Then, we took the best hyperparameters and trained models on data before the final timepoint, and evaluated on the final timepoint.

**Table 2** DeepEVFI Hyperparameters

| | C | D | F | Y-Ab | M-MP |
|---|---|---|---|---|---|
| MLP | 2x256 | 2x256 | 6x256 | 2x256 | 2x128 |
| Dropout | 20% | 20% | 0% | 0% | 0% |
| End-to-end loss | Dirichlet-multinomial | Dirichlet-multinomial | Multinomial | Multinomial | Multinomial |
| Weight decay | 1e-4 | 1e-4 | 1e-4 | 0 | 0 |
| Warmup | Yes | Yes | Yes | No | Yes |

All models used batch norm. We avoid optimization issues caused by the non-identifiability property by not performing minibatch optimization over variants. However, not all variants may be present in any single timepoint. In practice, we accumulate gradients and take one step per epoch.
